# Systematic identification of circular RNAs and corresponding regulatory networks unveil their potential roles in the midgut of *Apis cerana cerana* workers

**DOI:** 10.1101/558049

**Authors:** Dafu Chen, Huazhi Chen, Yu Du, Sihai Geng, Cuiling Xiong, Yanzhen Zheng, Chunsheng Hou, Qingyun Diao, Rui Guo

**Author notes:** These authors contributed equally to this work (Dafu Chen, Huazhi Chen).

## Abstract

**Background:** Circular RNAs (circRNAs) are newly discovered noncoding RNAs (ncRNAs) that play key roles in various biological functions, such as the regulation of gene expression and alternative splicing. CircRNAs have been identified in some species, including western honeybees. However, the understanding of honeybee circRNA is still very limited, and to date, no study on eastern honeybee circRNA has been conducted. Here, the circRNAs in the midguts of *Apis cerana cerana* workers were identified and validated, and the regulatory networks were constructed. Differentially expressed circRNAs (DEcircRNAs) and the corresponding competitively endogenous RNA (ceRNA) networks in the development of the worker’s midgut were further investigated.

**Results:** Here, 7- and 10-day-old *A. c. cerana* workers’ midguts (Ac1 and Ac2) were sequenced using RNA-seq, and a total of 9589 circRNAs were predicted using bioinformatics. These circRNAs were approximately 201-800 nt in length and could be classified into six types; the annotated exonic circRNAs were the most abundant. Additionally, five novel *A. c. cerana* circRNAs were confirmed by PCR amplification and Sanger sequencing, indicating the authenticity of *A. c. cerana* circRNAs. Interestingly, novel_circ_003723, novel_circ_002714, novel_circ_002451 and novel_circ_001980 were the most highly expressed circRNAs in both Ac1 and Ac2, which is indicative of their key roles in the development of the midgut. Moreover, 55 DEcircRNAs were identified in the Ac1 vs Ac2 comparison group, including 34 upregulated and 21 downregulated circRNAs. Further investigation showed that the source genes of circRNAs were classified into 34 GO terms and were involved in 141 KEGG pathways. In addition, the source genes of DEcircRNAs were categorized into 10 GO terms and 15 KEGG pathways, which demonstrated that the corresponding DEcircRNAs may affect the growth, development, and material and energy metabolisms of the worker’s midgut by regulating the expression of the related source genes. Additionally, the circRNA-miRNA regulatory networks were constructed and analyzed, and the results demonstrated that 1060 circRNAs can bind to 74 miRNAs and that 71.51% of circRNAs can be linked to only one miRNA. Furthermore, the DEcircRNA-miRNA-mRNA networks were constructed and explored, and the results indicate that the 13 downregulated circRNAs can bind to eight miRNAs and to 29 target genes. In addition, the results indicate that the 16 upregulated circRNAs can bind to 9 miRNAs and to 29 target genes, demonstrating that DEcircRNAs are likely involved in the regulation of midgut development via ceRNA mechanisms. Moreover, the regulatory networks of miR-6001-y-targeted DEcircRNAs were analyzed, and the results showed that eight DEcircRNAs may affect the development of *A. c. cerana* workers’ midguts by targeting miR-6001-y. Finally, four randomly selected DEcircRNAs were verified via RT-qPCR, confirming the reliability of our sequencing data.

**Conclusion:** This is the first systematic investigation of circRNAs and their corresponding regulatory networks in eastern honeybees. The identified circRNAs from the *A. c. cerana* worker’s midgut will enrich the known reservoir of honeybee ncRNAs. DEcircRNAs may play a comprehensive role during the development of the worker’s midgut via the regulation of source genes and the interaction with miRNAs by acting as ceRNAs. The eight DEcircRNAs that targeted miR-6001-y were likely to be vital for the development of the worker’s midgut. Our results provide a valuable resource for the future studies of *A. c. cerana* circRNA and lay a foundation to reveal the molecular mechanisms underlying the regulatory networks of circRNAs responsible for the worker’s midgut development; in addition, these findings facilitate a functional study on the key circRNAs involved in the developmental process.

**Graphical Abstract:** 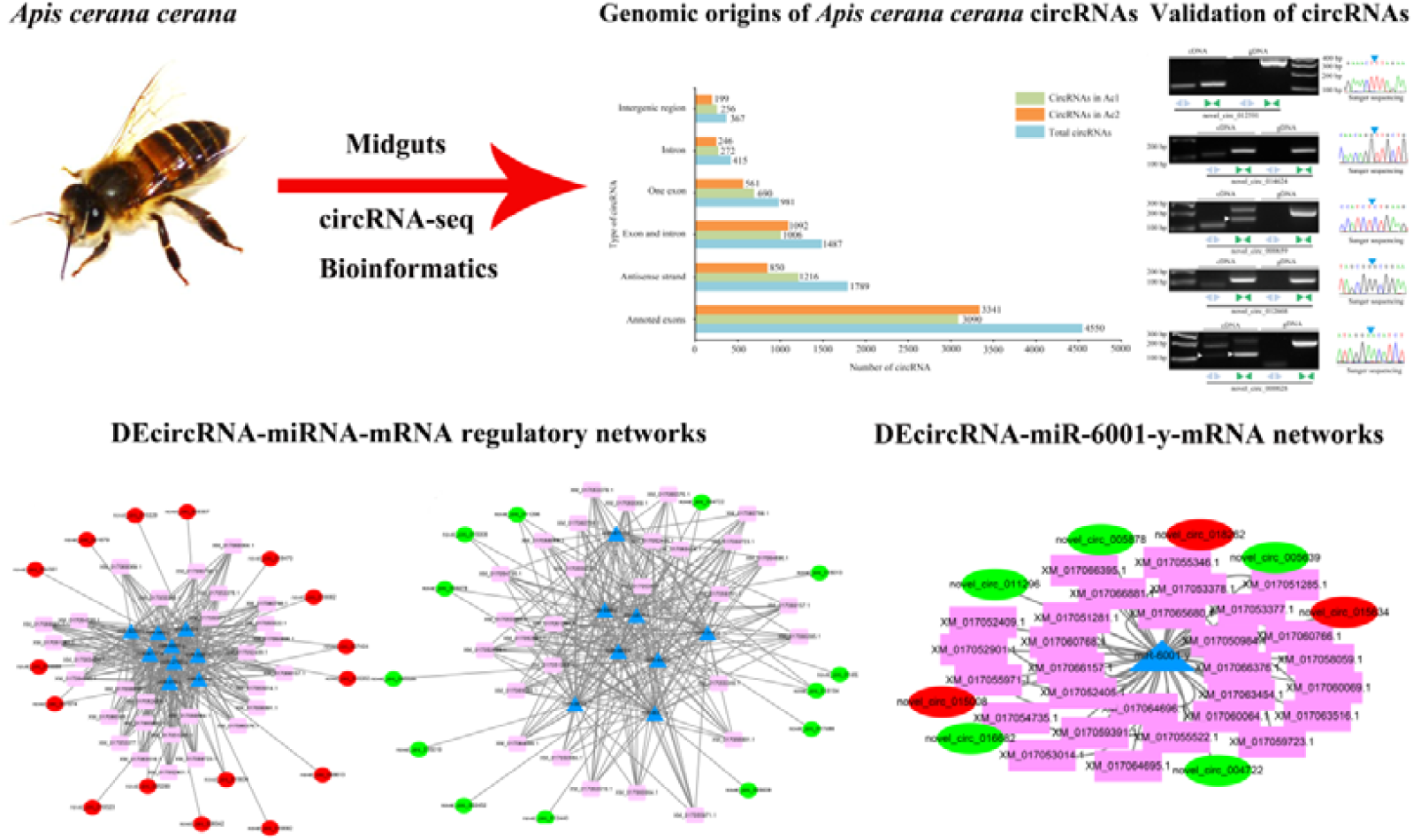

## Background

Circular RNAs (circRNAs), a newly discovered class of noncoding RNAs (ncRNAs), are lariated, intron paired, RNA-binding proteins (RBPs) or transcription factors formed into circular molecules [1, 2]. CircRNAs are conserved, abundant, and stable, and they have tissue- and temporal-specific expression in various species [3]. Previous studies have demonstrated that circRNAs were able to function as microRNA (miRNA) sponges, bound to RBPs and regulated the transcription of target genes and alternative splicing [4]. Recent studies have suggested that the circRNAs that contain ribosome entry sites [5] are capable of efficiently translating proteins [6]. Compared with linear RNA, circRNA is more stable and more resistant to digestion by the exonuclease RNase R due to a lack of 5’ cap and 3’ tail structures; thus, circRNA can be used as an ideal biomarker [7]. With the rapid development and application of high-throughput sequencing technologies and bioinformatic approaches, a number of circRNAs have been identified in humans [1, 8, 9], animals [10–15], plants [16–18], and microorganisms [19–23]. For example, Salzman et al. [9] predicted 46866 circRNAs in 15 human cell types and found that these circRNAs have potential regulatory functions. Shen and colleagues performed deep sequencing of different tissues from zebrafish, including the muscles, ovaries, and eyes, and identified a total of 3868 circRNAs based on the three following algorithms: find_circ, CIRI, and Segemehl. The authors further verified the expression of 84% of 176 circRNAs with high confidence [13]. Lu’s research group predicted 2354 circRNAs from rice using next-generation sequencing and further revealed that these circRNAs had a considerable number of isoforms [16]. Guo et al. [21] conducted high-throughput sequencing of mycelia and spore samples of *Ascosphaera apis* and discovered 551 circRNAs with a length of 200-600 nt using bioinformatics analysis; additionally, the authors investigated the regulatory networks between circRNAs and miRNAs and found complex interactions among them. Thus far, there is little research on insect circRNA, and the knowledge of circRNAs in insects, including honeybees, is still lacking [24–26]. Westholm et al. [24] analyzed the circRNAs of *Drosophila melanogaster* at the genome-wide level and found that the circRNA expression was tissue-specific. Recently, 3155 circRNAs were predicted from 1727 genes in the middle silk gland (MSG) and the posterior silk gland (PSG) of *Bombyx mori* by deep sequencing and bioinformatics analysis. Further analysis showed that the function and annotation information of the metabolic pathways of the source genes were similar. More recently, Chen et al. [26] predicted 12211 circRNAs from the ovarian tissue of *Apis mellifera ligustica* by a comparative analysis of virgin queens, egg-laying queens, egg-laying inhibited queens and egg-laying recovery queens; moreover, the authors speculated that the differentially expressed circRNAs (DEcircRNAs) played a role in the activation and spawning process of the ovarian tissue from the queen by competitively binding to miRNAs. It was reported that circRNAs function as miRNA sponges that naturally sequester and competitively suppress miRNA activity to regulate target gene expression [27, 28]. Yang’s group [28] performed a microarray analysis of human brain microvascular endothelial cells treated with methamphetamine and discovered that circHECW2 can competitively link to miR-30d as competitive endogenous RNA (ceRNA), thereby promoting the expression of the *autophagy-related 5* gene. A research group [29] analyzed the nucleus pulposus cells and tissues of patients with degenerative disc disease and found that circVMA21 could inhibit the apoptosis-related X protein-coding gene by adsorbing miR-200c. Hu and colleagues [30] analyzed the normal midgut tissue and the *B. mori* cytoplasmic polyhedrosis virus (BmCPV)-infected midgut tissue using circRNA-seq and bioinformatics analysis, and they respectively detected 9753 and 7475 circRNAs; the authors observed 294 upregulated and 106 downregulated circRNAs by comparative analysis, and they found that the alternative circularization of circRNAs was a common feature in silkworms and that the junction sites of many silkworm circRNAs were flanked by canonical GT/AG splicing signals.

The honeybee is the most important pollinator in nature and has irreplaceable economic and ecological value [31]; the honeybee is also a model insect for investigating development, host-pathogen interactions, social behavior and population genetics [32–34]. *Apis cerana cerana*, a subspecies of *Apis cerana*, is a specific bee species in China. It has the characteristics of being cold-tolerant, resistant to parasites, and good at collecting sporadic nectar sources [35]. The insect midgut plays a primary role in digesting food, absorbing nutrients and defending against pathogens [36, 37]. Previous studies have mainly concentrated on microorganisms in the bee gut [38, 39]. However, studies on the molecular mechanisms regulating the development of the honeybee gut are scarce, and the role of ncRNAs, including circRNAs, during the development of the honeybee gut is still unknown. In this study, we investigated the characteristics and the expression patterns of circRNAs in the midgut of eastern honeybees. We also explored the function of DEcircRNAs during the midgut development process by sequencing the midguts of *A. c. cerana* 7- and 10-day-old workers (Ac1 and Ac2) with high-throughput sequencing technology. This was followed by the identification and comparative analysis of circRNAs with bioinformatic approaches; in addition, randomly selected novel circRNAs were validated using conventional PCR with convergent and divergent primers. Furthermore, ceRNA regulatory networks of DEcircRNAs were constructed and investigated. Finally, DEcircRNAs were verified via real-time quantitative PCR (RT-qPCR). To our knowledge, this is the first documentation of the number, properties, expression patterns, differential expression profiles and potential biological functions of circRNAs in eastern honeybees.

## Results

### Overview of high-throughput sequencing data

On average, 124 and 99 million raw reads were generated from the Ac1 and Ac2 sample groups, respectively (**Table 1**). After filtering out the adapter sequences and the low-quality reads, 121 and 97 million clean reads with the Q30 means of 98.61% and 98.69%, respectively, were obtained (**Table 1**). In addition, Pearson correlations between the different biological repeats within the Ac1 and Ac2 groups were above 0.990 (Additional file 1). Together, these results confirmed that the samples and the sequencing data in this study were reasonable and reliable. Furthermore, the anchor reads of each sample were above 33360700, and the mapped anchors to the reference genome were above 27545100 (43.20%) (**Table 1**).

**Table 1.**
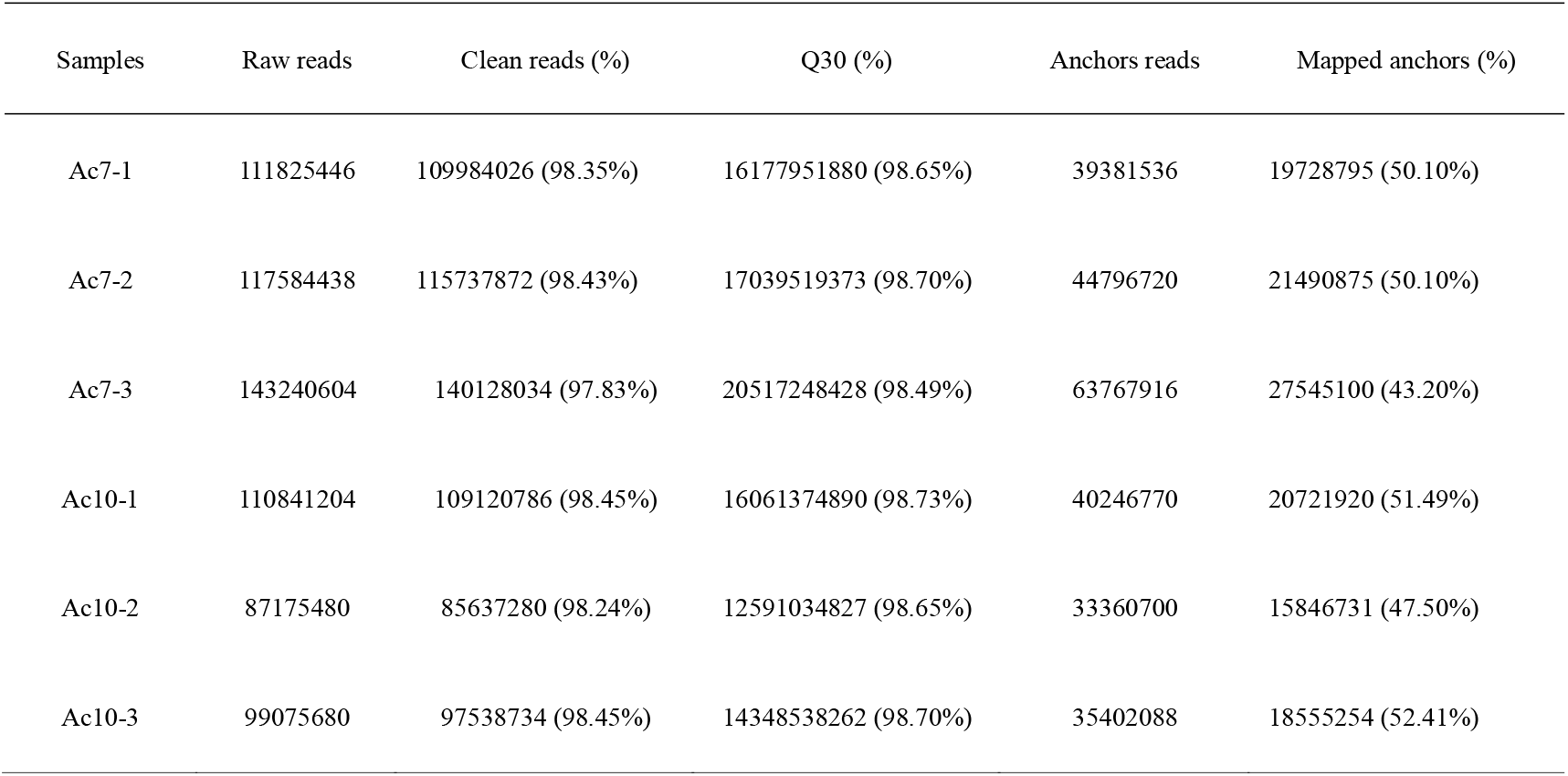
Overview of high-throughput sequencing datasets

### Identification and confirmation of circRNAs in *A. c. cerana* workers’ midguts

Here, a total of 9589 circRNAs were predicted from the midguts of *A. c. cerana* workers using bioinformatics software (**Fig. 1**). Among them, 3230 (33.7%) circRNAs were shared by Ac1 and Ac2, and the respective numbers of unique circRNAs were 3300 (34.4%) and 3059 (31.9%) (**Fig. 2a**). Additionally, the length of the circRNAs mainly ranged from 201 nt to 800 nt; the shortest circRNA was 15 nt, and the longest circRNA was 84596 nt (**Fig. 2b**). Further analysis showed that 3.8%, 4.3%, 10.2%, 15.5%, 18.7%, and 47.5% of circRNAs were intergenic, intronic, single exon, intron-exon, antisense, and annotated-exons (multiple exons), respectively (**Fig. 2c**).

**Fig. 1.**
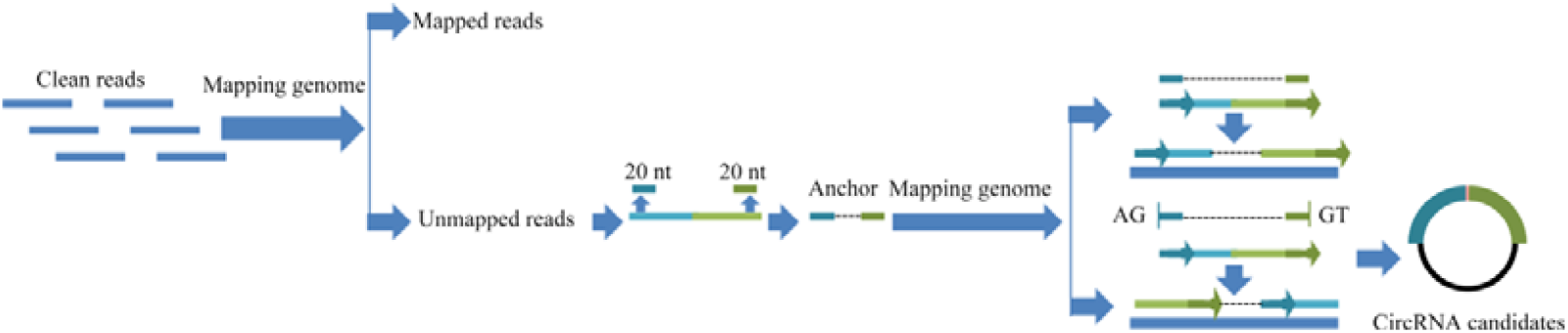
Computational pipeline for the prediction of circRNAs in the midguts of *A. c. cerana* workers.

**Fig. 2.**
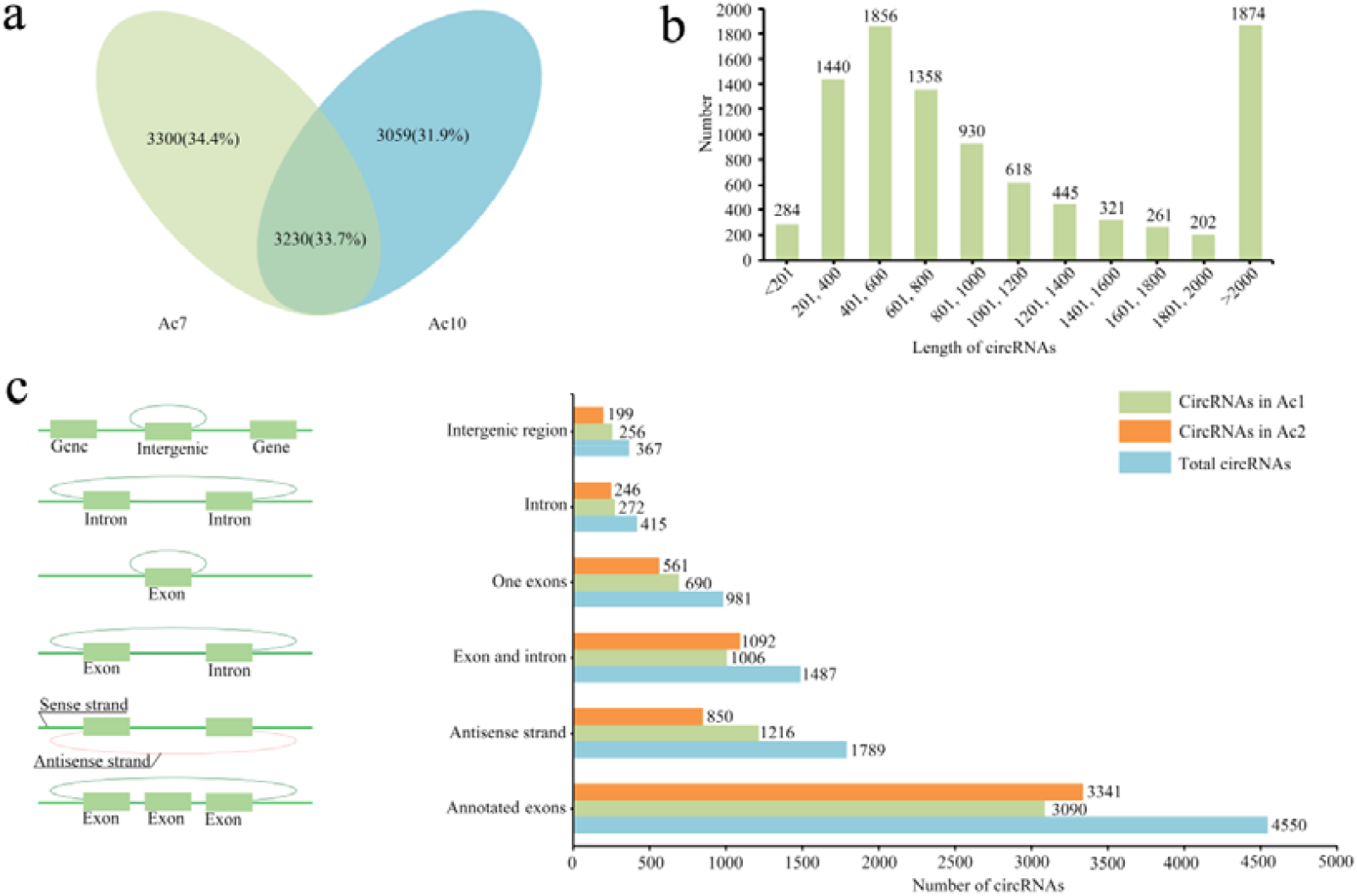
Structural characteristics of circRNAs in the midguts of *A. c. cerana* workers. **a** A Venn diagram of circRNAs in Ac1 and Ac2. **b** The length distribution of circRNAs. **c** The genomic origin of circRNAs; Left: Six types of source genes of circRNAs. Right: The number of circRNAs from various types of source genes.

To experimentally confirm the candidate *A. c. cerana* circRNAs, convergent and divergent primers (**Fig. 3a**; Additional file 2: **Table S1**) were designed to amplify each circRNA using both the RNase R-treated sample (cDNA) and the genomic DNA (gDNA) as PCR templates. Due to the circular structure, circRNA is more resistant to digestion by the exonuclease RNase R than linear RNA is (**Fig. 3b**). Theoretically, convergent primers should amplify products from both the cDNA and the gDNA, but the divergent primers should only amplify the circRNAs from the cDNA. Distinct PCR products with the expected size were amplified using the convergent and divergent primers, and the back-splicing sites were verified by Sanger sequencing (**Fig. 3c**). These results indicated that the predicted *A. c. cerana* circRNAs in the current study were credible.

**Fig. 3.**
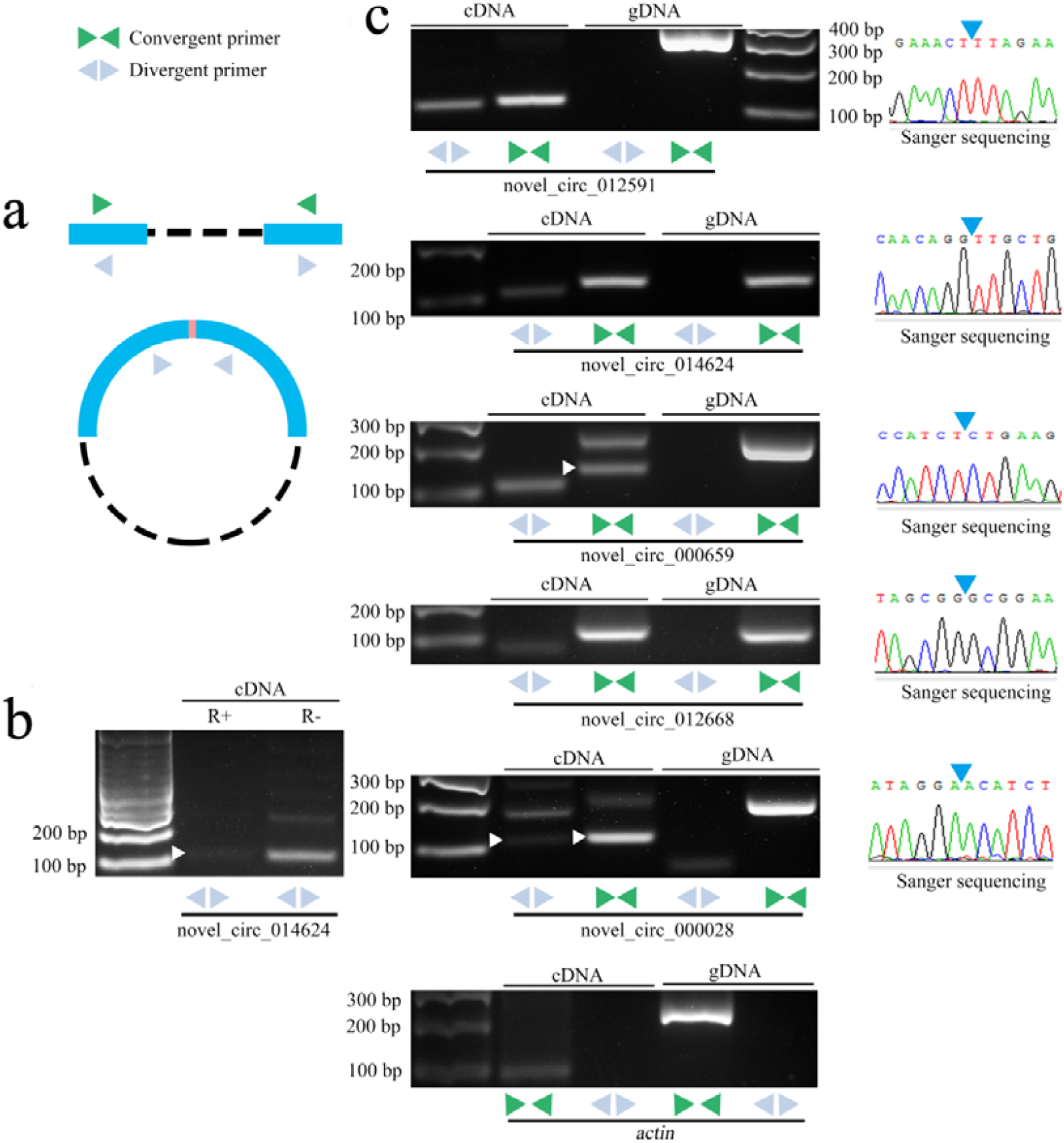
Validation of *A. c. cerana* circRNAs via PCR using convergent and divergent primers. **a** A schematic diagram showing the strategy of circRNA validation. Convergent primers were designed to detect linear RNA, and divergent primers were designed to detect circRNAs. **b** *A. c. cerana* circRNA (novel_circ_014624) was amplified with divergent primers using RNase R-digested RNA and undigested RNA as templates. “R+” indicates RNA treated with RNase R; “R−” indicates untreated RNA. **c** The agarose gel electrophoresis (left) and Sanger sequencing (right) of RT-PCR products is shown. Divergent primers (gray back-to-back triangle pairs) successfully amplified five circRNAs (novel_circ_012591, novel_circ_014624, novel_circ_000659, novel_circ_012668 and novel_circ_000028) in cDNA but failed to do so in gDNA. Convergent primers (green opposing triangle pairs) worked on both cDNA and gDNA. *A. c. cerana actin* was used as a linear control, and Sanger sequencing further confirmed the head-to-tail back-splicing sites of circRNAs (blue triangles).

### Expression profiles of circRNAs in the midgut of an *A. c. cerana* worker

The RPM value for each circRNA was calculated and analyzed, and the results suggested that novel_circ_003723 (RPM=123156.9), novel_circ_002714 (RPM=25893.1), novel_circ_002451 (RPM=23955.74), novel_circ_001980 (RPM=23566.74), and novel_circ_002595 (RPM=17669.37) were the highest expressed circRNAs in Ac1 (**Table 2**); however, novel_circ_003723 (RPM=115167.7), novel_circ_002714 (RPM=29018.24), novel_circ_002451 (RPM=28794.78), novel_circ_001980 (RPM=22996.79), and novel_circ_007006 (RPM=16983.2) were the highest expressed circRNAs in Ac2 (**Table 3**). Interestingly, nine were shared by Ac1 and Ac2, implying that they have important roles during the development of the *A. c. cerana* worker’s midgut. In addition, novel_circ_006624 (RPM=11006.91) and novel_circ_002101 (RPM=11422.22) with high expression level were specific in Ac1 and Ac2, which indicated that these two circRNAs were likely to play special roles in different stages of midgut development.

**Table 2.** Top 10 expressed circRNAs in Ac1 sample group

| CircRNA | RPM | Source gene ID | Length | Type of circularization |
| --- | --- | --- | --- | --- |
| novel_circ_003723 | 123156.9 | 107992487 | 552 | annotated exons |
| novel_circ_002714 | 25893.1 | 108004068 | 10897 | exon intron |
| novel_circ_002451 | 23955.74 | 108004042 | 1190 | one exon |
| novel_circ_001980 | 23566.74 | 108003484 | 528 | annotated exons |
| novel_circ_002595 | 17669.37 | 108003973 | 1043 | annotated exons |
| novel_circ_007006 | 14298.63 | 107994933 | 1742 | annotated exons |
| novel_circ_014464 | 14249.72 | 107999817 | 3776 | exon intron |
| novel_circ_006624 | 11006.91 | 107994327 | 705 | annotated exons |
| novel_circ_017811 | 10093.41 | 108002419 | 1818 | exon intron |
| novel_circ_015368 | 9880.122 | 108000581 | 8118 | exon intron |

**Table 3.** Top 10 expressed circRNAs in Ac2 sample group

| CircRNA | RPM | Source gene ID | Length | Type of circularization |
| --- | --- | --- | --- | --- |
| novel_circ_003723 | 115167.7 | 107992487 | 552 | annotated exons |
| novel_circ_002714 | 29018.24 | 108004068 | 10897 | exon intron |
| novel_circ_002451 | 28794.78 | 108004042 | 1190 | one exon |
| novel_circ_001980 | 22996.79 | 108003484 | 528 | annotated exons |
| novel_circ_007006 | 16983.2 | 107994933 | 1742 | annotated exons |
| novel_circ_002595 | 15959.88 | 108003973 | 1043 | annotated exons |
| novel_circ_014464 | 14755.94 | 107999817 | 3776 | exon intron |
| novel_circ_002101 | 11422.22 | 108003728 | 6735 | exon intron |
| novel_circ_017811 | 9971.993 | 108002419 | 1818 | exon intron |
| novel_circ_015368 | 9861.83 | 108000581 | 8118 | exon intron |

### Differential expression of circRNAs in the development of the *A. c. cerana* worker’s midgut

In the Ac1 vs Ac2 comparison group, there were 55 DEcircRNAs, including 34 upregulated and 21 downregulated circRNAs (**Fig. 4a**). The heat map indicated that these DEcircRNAs had various expression levels (**Fig. 4b**) as follows: the mostly upregulated circRNAs were novel_circ_005299 (log_2_ fold change=16.95329), novel_circ_015634 (log_2_ fold change=16.72073), novel_circ_001191 (log_2_ fold change=16.46338) (**Table 4**); however, novel_circ_018184 (log_2_ fold change=-17.0462), novel_circ_012440 (log_2_ fold change=-17.0158), and novel_circ_004722 (log_2_ fold change=-16.7506) were the DEcircRNAs with the highest downregulation (**Table 4**).

**Fig. 4.**
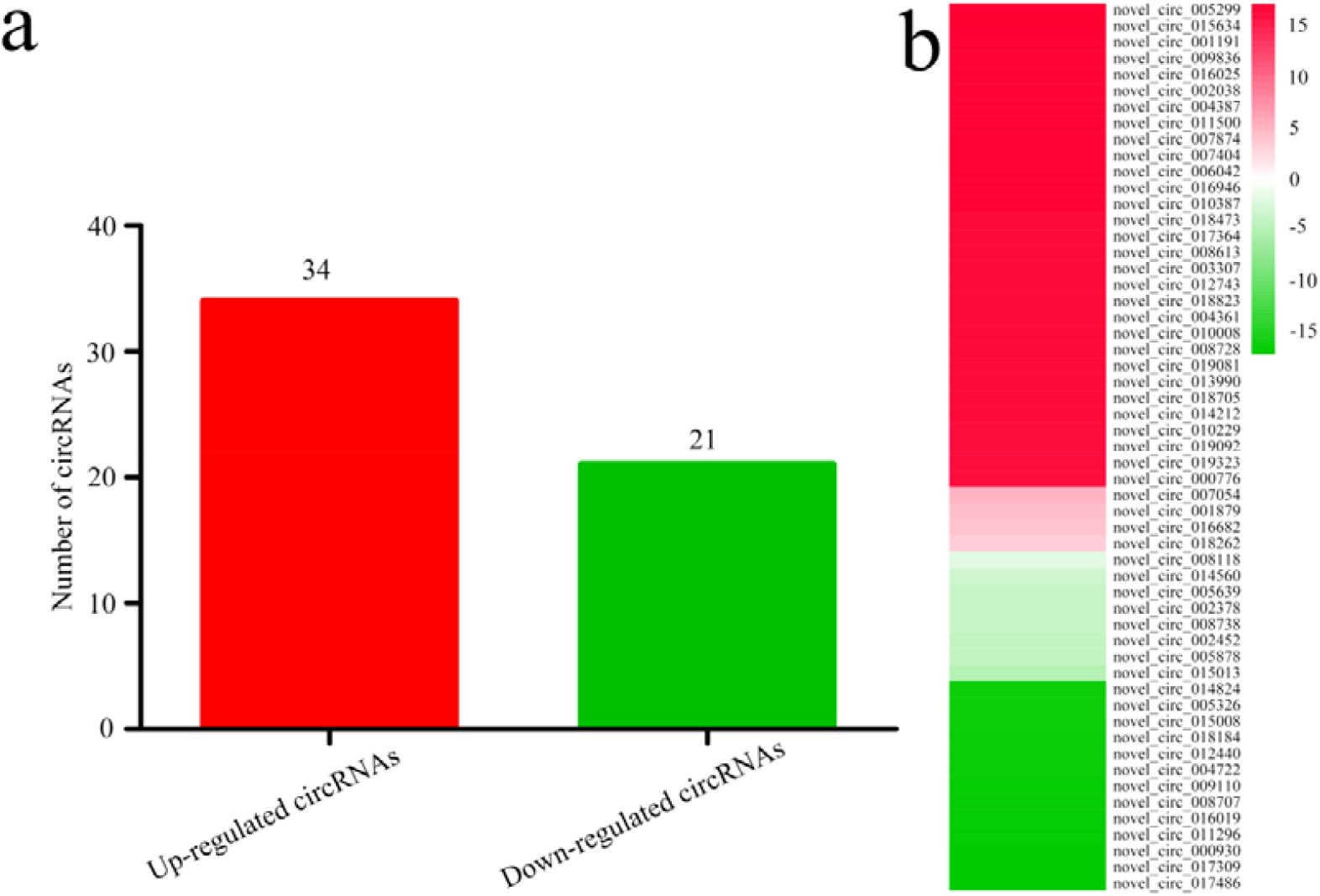
Differential expression profiles of DEcircRNAs in the Ac1 vs Ac2 comparison group. **a** The number of up- and downregulated circRNAs. **b** The expression clustering of DEcircRNAs.

**Table 4.** Detailed information of DEcircRNAs in Ac1 vs Ac2 comparison group.

| cicRNA | Log2(Fold<br>change) | RPM value in Ac7 | RPM value in Ac10 | P value | Source gene ID |
| --- | --- | --- | --- | --- | --- |
| novel_circ_005299 | 16.95329 | 0.001 | 126.8965 | 0.001168 | 107993571 |
| novel_circ_015634 | 16.72073 | 0.001 | 108.0046 | 0.002029 | 108000772 |
| novel_circ_001191 | 16.46338 | 0.001 | 90.35904 | 0.005439 | 108001166 |
| novel_circ_009836 | 16.44963 | 0.001 | 89.50156 | 0.023023 | 107996840 |
| novel_circ_016025 | 16.44047 | 0.001 | 88.93513 | 0.00749 | 108000985 |
| novel_circ_002038 | 16.43654 | 0.001 | 88.69351 | 0.005915 | 108003721 |
| novel_circ_004387 | 16.41845 | 0.001 | 87.5882 | 0.012274 | 107992925 |
| novel_circ_011500 | 16.38751 | 0.001 | 85.72992 | 0.026449 | 107998190 |
| novel_circ_007874 | 16.33445 | 0.001 | 82.63411 | 0.007655 | 107995296 |
| novel_circ_007404 | 16.3249 | 0.001 | 82.08902 | 0.008573 | 107995050 |
| novel_circ_006042 | 16.32095 | 0.001 | 81.86436 | 0.010107 | 107994002 |
| novel_circ_016946 | 16.30991 | 0.001 | 81.2403 | 0.033485 | 108001789 |
| novel_circ_010387 | 16.29053 | 0.001 | 80.15633 | 0.038359 | 107997309 |
| novel_circ_018473 | 16.26564 | 0.001 | 78.7855 | 0.037914 | 108002943 |
| novel_circ_017364 | 16.25641 | 0.001 | 78.28291 | 0.042661 | 108002081 |
| novel_circ_008613 | 16.20899 | 0.001 | 75.7517 | 0.012921 | 107995818 |
| novel_circ_003307 | 16.18952 | 0.001 | 74.73612 | 0.013273 | 108004261 |
| novel_circ_012743 | 16.16319 | 0.001 | 73.3848 | 0.013852 | 107998821 |
| novel_circ_018823 | 16.12723 | 0.001 | 71.57813 | 0.018403 | 108002989 |
| novel_circ_004361 | 16.11451 | 0.001 | 70.94952 | 0.017185 | 107992916 |
| novel_circ_010008 | 16.08121 | 0.001 | 69.33104 | 0.019425 | 107996875 |
| novel_circ_008728 | 16.01634 | 0.001 | 66.28228 | 0.025863 | 107996102 |
| novel_circ_019081 | 15.98466 | 0.001 | 64.84307 | 0.044915 | 108003259 |
| novel_circ_013990 | 15.97469 | 0.001 | 64.39646 | 0.026724 | 107999331 |
| novel_circ_018705 | 15.96411 | 0.001 | 63.92596 | 0.029844 | 108003021 |
| novel_circ_014212 | 15.96151 | 0.001 | 63.81053 | 0.027349 | 107999607 |
| novel_circ_010229 | 15.92102 | 0.001 | 62.04469 | 0.047502 | 107997067 |
| novel_circ_019092 | 15.8814 | 0.001 | 60.36403 | 0.036297 | 108003256 |
| novel_circ_019323 | 15.86527 | 0.001 | 59.69275 | 0.046722 | 108003377 |
| novel_circ_000776 | 15.81098 | 0.001 | 57.48834 | 0.03993 | 107997219 |
| novel_circ_007054 | 5.024941 | 4.421717 | 143.9623 | 0.004653 | 107994823 |
| novel_circ_001879 | 4.231173 | 4.421717 | 83.0426 | 0.042954 | 108003674 |
| novel_circ_016682 | 3.964923 | 13.26515 | 207.1442 | 0.018444 | 108001529 |
| novel_circ_018262 | 3.297969 | 12.94539 | 127.3218 | 0.042754 | 108002803 |
| novel_circ_017486 | -17.0462 | 135.3374 | 0.001 | 0.000928 | 108002060 |
| novel_circ_017309 | -17.0158 | 132.518 | 0.001 | 0.008704 | 108002079 |
| novel_circ_000930 | -16.7506 | 110.2628 | 0.001 | 0.001198 | 107999008 |
| novel_circ_011296 | -16.5499 | 95.94388 | 0.001 | 0.018969 | NA |
| novel_circ_016019 | -16.5071 | 93.13666 | 0.001 | 0.027868 | 108001004 |
| novel_circ_008707 | -16.4751 | 91.09608 | 0.001 | 0.022043 | 107996048 |
| novel_circ_009110 | -16.4196 | 87.65767 | 0.001 | 0.02767 | 107996165 |
| novel_circ_004722 | -16.2595 | 78.4521 | 0.001 | 0.03955 | 107993104 |
| novel_circ_012440 | -16.2365 | 77.21265 | 0.001 | 0.039603 | 107998618 |
| novel_circ_018184 | -16.1698 | 73.7198 | 0.001 | 0.047717 | 108002833 |
| novel_circ_015008 | -15.9373 | 62.75077 | 0.001 | 0.030674 | 108000196 |
| novel_circ_005326 | -15.8361 | 58.49951 | 0.001 | 0.041909 | 107993584 |
| novel_circ_014824 | -15.8064 | 57.30567 | 0.001 | 0.047929 | 108000053 |
| novel_circ_015013 | -4.9629 | 73.01616 | 2.34118567 | 0.048552 | NA |
| novel_circ_005878 | -3.91784 | 141.5411 | 9.36474268 | 0.03115 | 107993914 |
| novel_circ_002452 | -3.88818 | 103.9958 | 7.02355701 | 0.029129 | 108004042 |
| novel_circ_008738 | -3.73248 | 124.4754 | 9.36474268 | 0.041704 | 107996056 |
| novel_circ_002378 | -3.60814 | 228.3945 | 18.72948536 | 0.037572 | 108003872 |
| novel_circ_005639 | -3.48506 | 138.1117 | 12.33453461 | 0.027156 | 107993779 |
| novel_circ_014560 | -3.33887 | 168.6312 | 16.66622071 | 0.016433 | 107999904 |
| novel_circ_008118 | -1.97648 | 421.6318 | 107.1406954 | 0.039033 | 107995510 |

### Putative functions of all circRNAs and DEcircRNAs in the midguts of *A. c. cerana* workers

Intronic circRNAs or exonic and intronic circRNAs were able to regulate the expression of source genes [40]. To explore the putative function of all predicted circRNAs and DEcircRNAs, GO term and KEGG pathway analyses for the source genes were conducted. The results demonstrated that the source genes for all of the circRNAs can be categorized into 34 GO terms (**Fig. 5a**; Additional file 3: **Table S2**). The top three GO terms for each subgroup are as follows: the terms for the cellular component (CC) subgroup were cell part, cell, and organelle; the terms for the biological process (BP) subgroup were cellular process, single-organism process, and metabolic process; and the terms for the molecular function (MF) subgroup were binding, catalytic activity, and molecular transducer activity. These results suggest that circRNAs play comprehensive roles in cell components, biological processes, and molecular functions in the midguts of *A. c. cerana* workers. Additionally, the source genes of DEcircRNAs were categorized into 10 GO terms as follows: catalytic activity, binding, molecular transducer activity, transporter activity, signal transducer activity, cellular process, metabolic process, cell part, cell, and organelle (**Fig. 5b**; Additional file 4: **Table S3**). This indicates that the regulation of source genes by DEcircRNAs for the regulation of midgut development is associated with various biological functions.

**Fig. 5.**
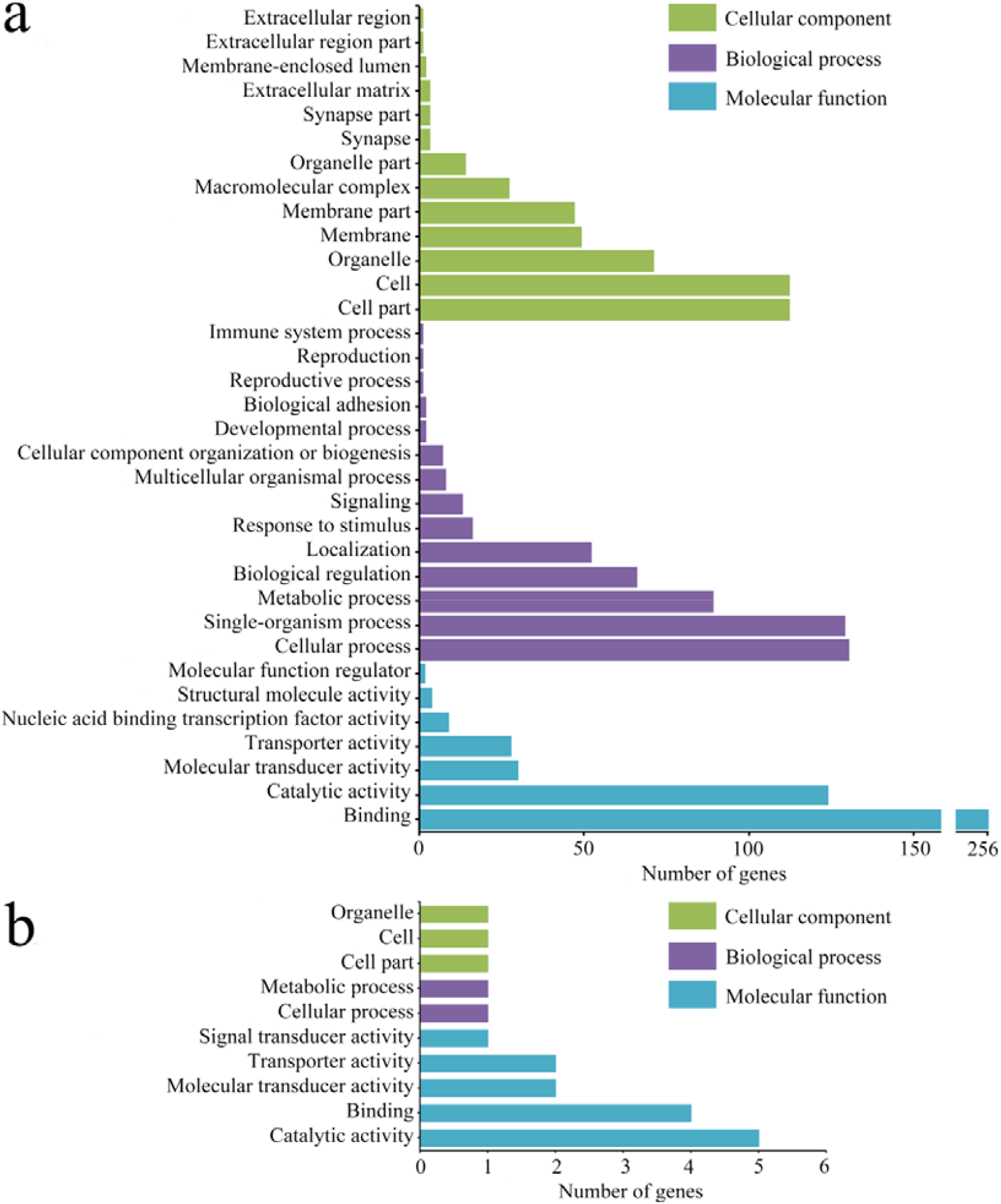
GO analyses for the source genes of total *A. c. cerana* circRNAs and DEcircRNAs in the Ac1 vs Ac2 comparison group. **a** Total *A. c. cerana* circRNAs. **b** DEcircRNAs in the Ac1 vs Ac2 group.

In addition, the source genes of circRNAs were engaged in as many as 141 pathways (Additional file 5: **Table S4**), including protein processing in the endoplasmic reticulum, endocytosis, the Wnt signaling pathway, the Hippo signaling pathway, and purine metabolism (**Fig. 6a**); this further suggests the general function of *A. c. cerana* circRNAs. The source genes of DEcircRNAs were involved in 15 pathways, including ABC transporters, sphingolipid metabolism, ECM-receptor interaction, glycerophospholipid metabolism, neuroactive ligand-receptor interaction, the Hippo signaling pathway, lysosome, pyruvate metabolism, citrate cycle (TCA cycle), glycolysis/gluconeogenesis, insulin resistance, the FoxO signaling pathway, ubiquitin-mediated proteolysis, carbon metabolism, and purine metabolism (**Fig. 6b**; Additional file 6: **Table S5**); this result indicated that DEcircRNAs were likely to play a regulatory role in some important pathways associated with signaling, immunity, and material and energy metabolism, thereby controlling the development of the midgut.

**Fig. 6.**
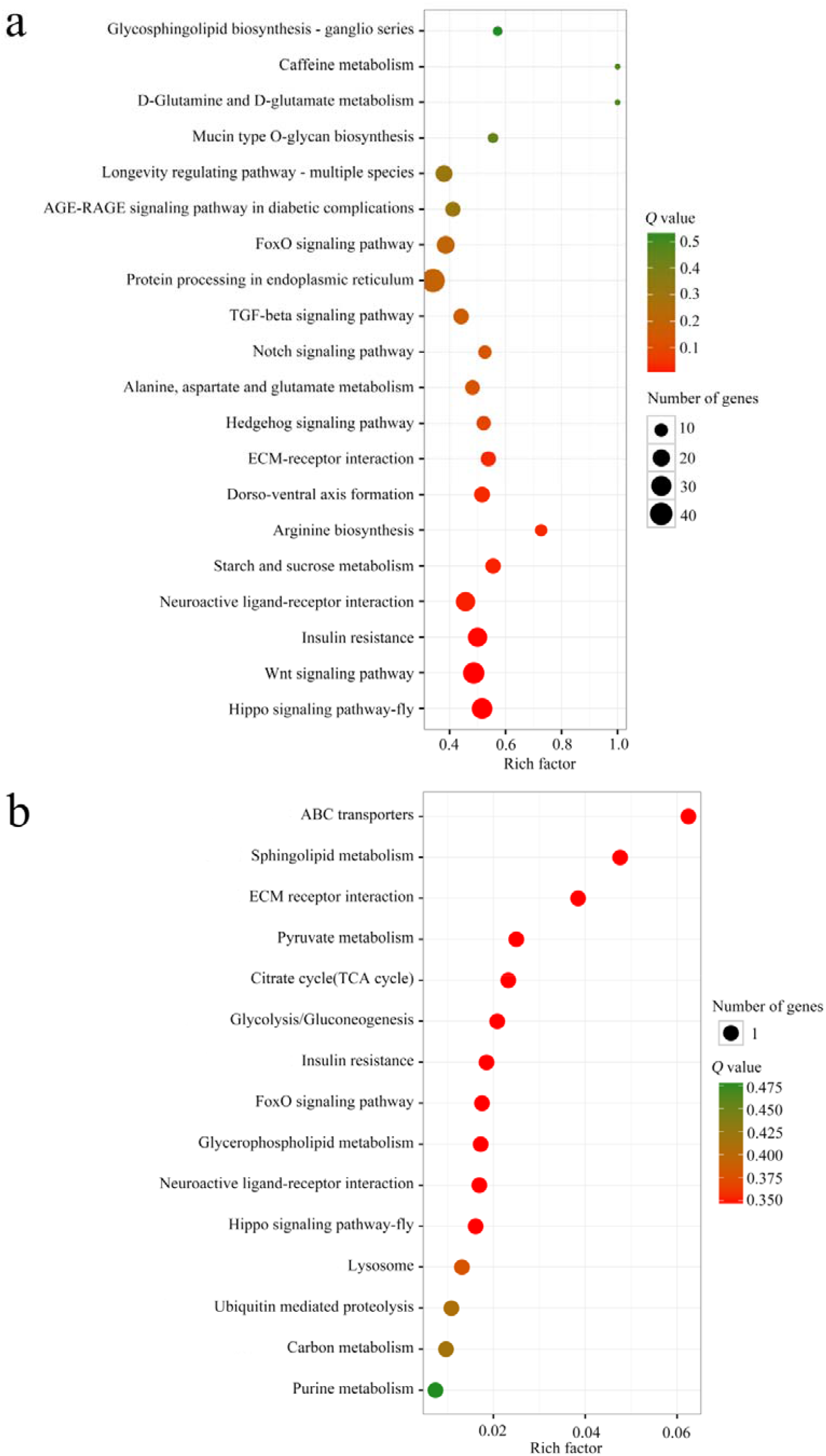
KEGG enrichment analyses for the source genes of total *A. c. cerana* circRNAs and DEcircRNAs in the Ac1 vs Ac2 comparison group. **a** Total *A. c. cerana* circRNAs. **b** DEcircRNAs in the Ac1 vs Ac2 group. The *x*-axis represents the rich factor (the ratio of input number and background number), which presents the degree of enrichment. Green and red respectively indicate high and low *Q* values; the greener, the higher the value, and the redder, the lower the value. The *y*-axis represents pathway terms. The size of the circle indicates the number of enriched genes in a certain pathway, the larger the circle, the higher the number.

### CircRNA-miRNA regulatory networks in the midguts of *A. c. cerana* workers

In the present study, the regulatory networks of total circRNAs and their corresponding target miRNAs were constructed and investigated (**Fig. 7a**; see also Additional file 7: **Table S6**). A total of 1060 circRNAs were found to be targeted by 74 miRNAs. Among them, 758 (71.51%) circRNAs were found to bind to only one miRNA, such as novel_circ_003086, novel_circ_003087 and novel_circ_007792. However, some circRNAs could bind to several target miRNAs. For instance, miR-4962-y, miR-6873-y, miR-1895-y and miR-265-y had 328, 294, 146 and 100 target miRNAs, respectively; in addition, novel_circ_004310, novel_circ_004312, novel_circ_012714 and novel_circ_012715 were found to bind to eight target miRNAs. The results indicate that complex circRNA-miRNA regulatory networks exist in the midgut of *A. c. cerana*, and circRNAs may function via indirectly regulating gene expression mediated by miRNAs. Moreover, we randomly selected 13 miRNAs in the networks for stem-loop RT-PCR, and the results showed that the majority of miRNAs (12/13) could be amplified, which proved the reality of miRNAs in the network (**Fig. 7b**; Additional file 8: **Table S7**).

**Fig. 7.**
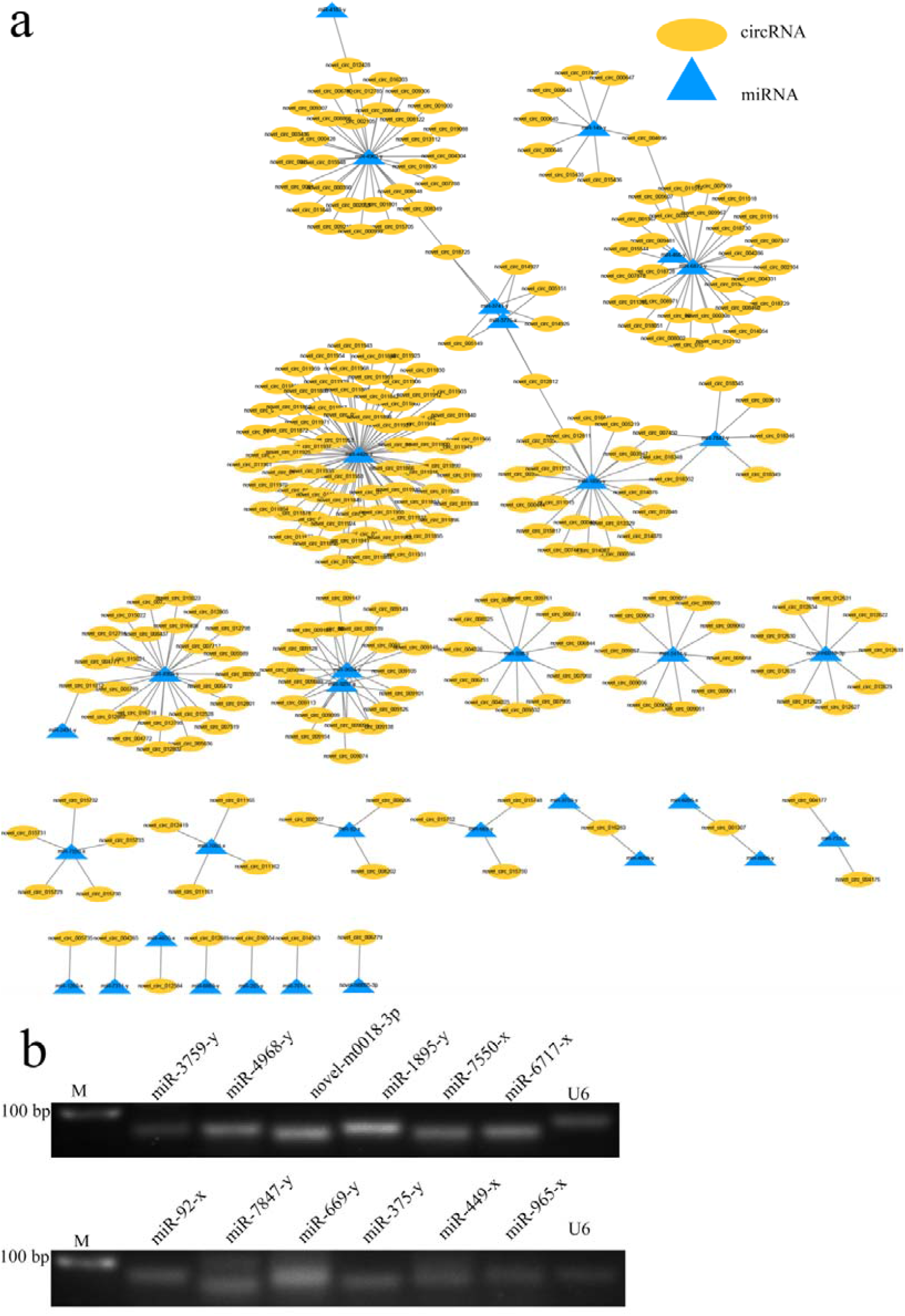
The circRNA-miRNA regulatory networks in the midguts of *A. c. cerana* workers. **a** Regulatory networks of the top 300 circRNAs and their target miRNAs. **b** The validation of circRNA target miRNAs in the center of regulatory networks using stem-loop RT-PCR.

### DEcircRNA-miRNA and DEcircRNA-miRNA-mRNA regulatory networks in the midguts of *A. c. cerana* workers

Recent evidence has shown that RNAs regulate each other with miRNA response elements (MREs) through a ceRNA mechanism [41–43]. To explore the ceRNA regulatory network of DEcircRNAs in the midgut, DEcircRNA-miRNA regulatory networks were constructed and analyzed (Additional file 9: **Table S8**). The results showed that 16 upregulated circRNAs linked to nine miRNAs, among which, novel_circ_015634 bound to three miRNAs, miR-598-y, miR-993-y and miR-6001-y (**Fig. 8a**), while 13 downregulated circRNAs bound to eight miRNAs; for example, novel_circ_011296 bound to miR-993-y, miR-6001-y and miR-315-x (**Fig. 8b**).

**Fig. 8.**
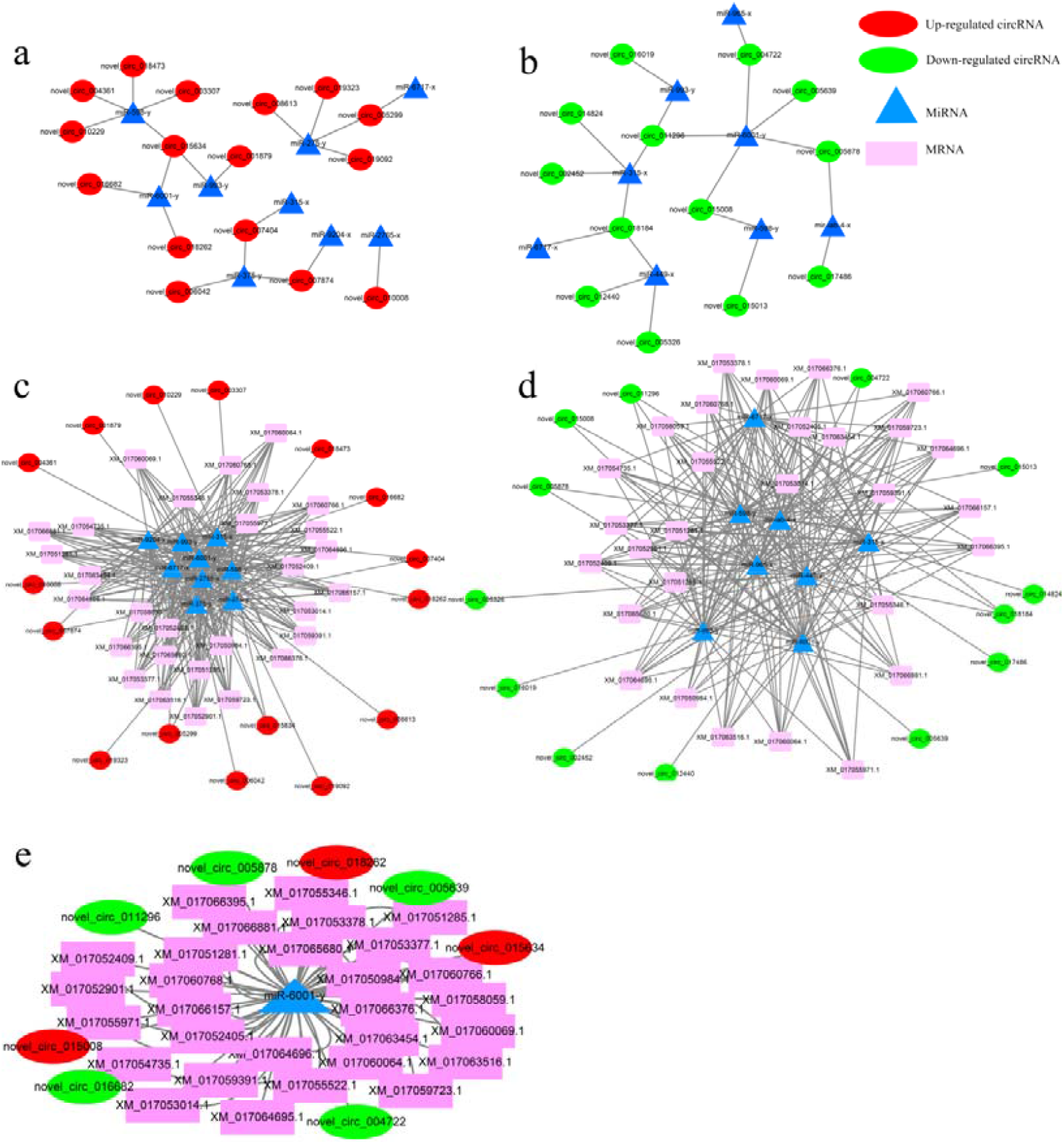
Regulatory networks of DEcircRNA in the midguts of *A. c. cerana* workers. **a** Regulatory networks of upregulated circRNA and their target miRNAs. **b** Regulatory networks of downregulated circRNA and their target miRNAs. **c** Regulatory networks of upregulated circRNAs, their target miRNAs and the miRNA-targeted mRNAs. **d** Regulatory networks of downregulated circRNA, their target miRNAs, and the miRNA-targeted mRNAs. **e** Regulatory networks of miR-6001-y, miR-6001-y-targeted DEcircRNAs and miR-6001-y-targeted mRNAs.

Furthermore, the regulatory networks of DEcircRNA-miRNA-mRNA were constructed to explore the potential role of DEcircRNAs (Additional file 10: **Table S9**). The results demonstrated that nine target miRNAs of upregulated circRNAs could bind to 29 mRNAs (**Fig. 8c**), while eight target miRNAs of downregulated circRNAs could bind to 29 mRNAs (**Fig. 8c**). These results indicate that there are complex cross-talks between DEcircRNAs, miRNAs and mRNAs and that DEcircRNAs are likely to participate in the regulation of the development of the midgut via ceRNA regulatory networks. The regulatory network of DEcircRNA-miR-6001-y-mRNA was presented in **Fig. 8e**.

### Validation of DEcircRNAs in the *A. c. cerana* worker’s midgut via RT-qPCR

To validate the deep sequencing data in this work, four randomly selected DEcircRNAs were analyzed using RT-qPCR. The results indicate that their expression levels are consistent with those detected using next-generation sequencing (**Fig. 9**; Additional file 11: **Table S10**), which validates the reliability of the sequencing data in this work.

**Fig. 9.**
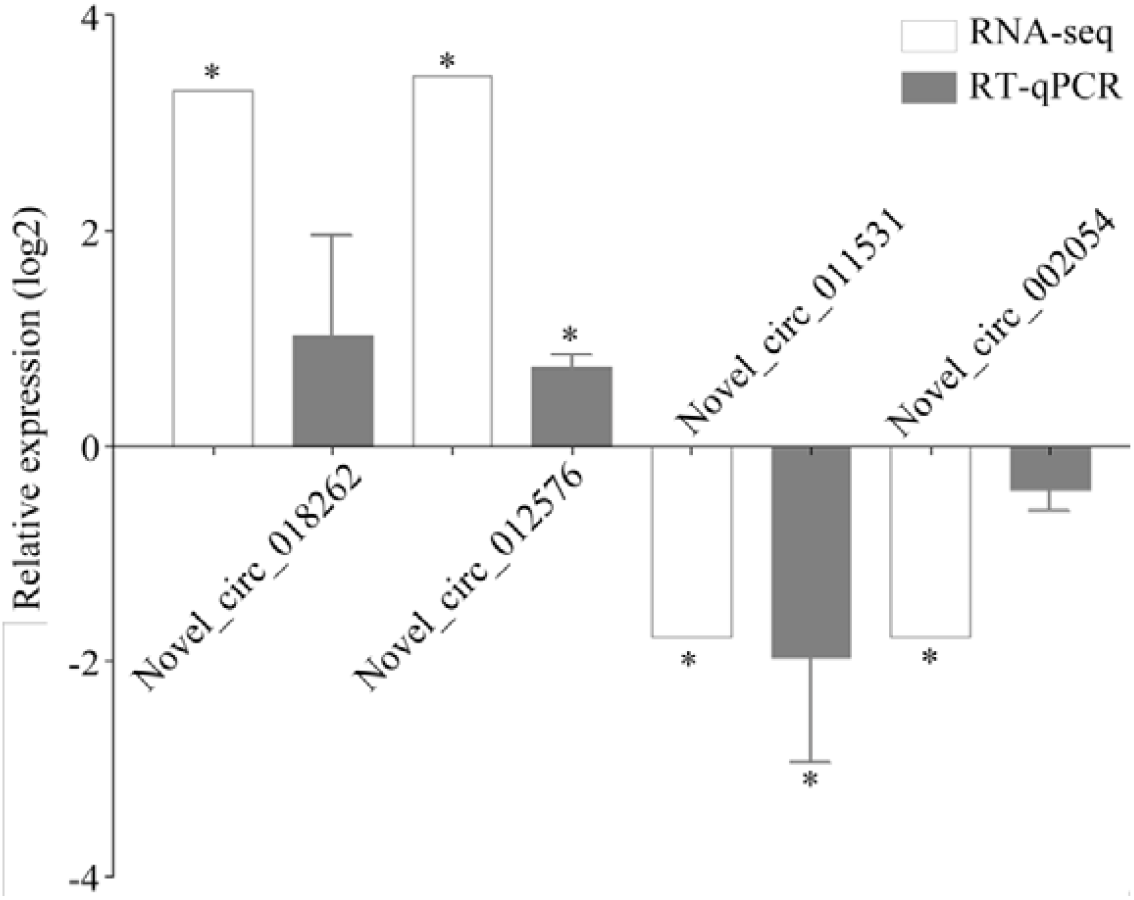
RT-qPCR confirmation of *A. c. cerana* DEcircRNAs. “*” represents *P* value < 0.05.

## Discussion

Recent evidence suggests that circRNAs are able to play very important roles in alternative splicing, transcriptional regulation, and the regulation of source gene expression. With the rapid development of high-throughput sequencing technologies and bioinformatics analyses, circRNAs have been identified in many species, including humans [1, 8, 9], mice [10], cattle [11], grass carp [12], zebrafish [13], chickens [14], pigs [15], rice [16], barley [17], *Arabidopsis thaliana* [18], Archaea [19], *Caenorhabditis elegans* [20], *A. apis* [21], *Nosema ceranae* [22], and Epstein-Barr virus [23]. In insects such as *D. melanogaster, B. mori* and *A. m. ligustica*, circRNAs have also been reported [24–26]. More recently, Chen’s research group detected 12211 circRNAs in the ovaries of the *A. m. ligustica* queen bee, and more than 80% of the circRNAs contained circular exons (one exon and annotated exons); the authors further observed that 1340, 175 and 100 circRNAs were differentially expressed when comparing the circRNAs of egg-laying queens and virgin queens, egg-laying inhibited queens and egg-laying queens, as well as egg-laying recovery queens and egg-laying inhibited queens [26]. To the best of our knowledge, there is no report of circRNA in eastern honeybees, including *A. c. cerana*. In the present study, 7- and 10-day-old *A. c. cerana* workers’ midguts were sequenced using the Illumina HiSeq platform, and based on the bioinformatic analysis, 6530 and 6359 circRNAs were predicted, respectively. In addition, 3230 circRNAs were shared by Ac1 and Ac2, and the respective numbers of unique circRNAs were 3300 (34.4%) and 3059 (31.9%). The length of *A. c. cerana* circRNAs was mainly 201~800 nt. In addition, these circRNAs could be divided into six types according to their location in the *A. cerana* genome, and exonic circRNAs (66.2%) were found to be the most abundant. This finding is similar to the *D. melanogaster* and *A. m. ligustica* circRNAs [24, 26] but is different from the *B. mori* circRNAs [25], and this demonstrated that circRNAs are abundant and species-specific. The identified *A. c. cerana* circRNAs will enrich the known reservoir of bee ncRNAs and will be a valuable resource for future research. To verify the back-splicing junction of the identified circRNAs, the expression and back-splicing junctions of four *A. c. cerana* circRNAs were experimentally verified by PCR amplification and Sanger sequencing (**Fig. 3c**).

In this study, the nine following *A. c. cerana* circRNAs: novel_circ_003723, novel_circ_002714, novel_circ_002451, novel_circ_001980, novel_circ_002595, novel_circ_007006, novel_circ_014464, novel_circ_017811, and novel_circ_015368, were found to be highly expressed in both Ac1 and Ac2, which is indicative of their crucial functions in the development of *A. c. cerana* worker’s midgut. However, whether these circRNAs are also highly expressed in the midgut during other developmental stages remains unknown and needs to be further studied. Interestingly, novel_circ_006624 (RPM=11006.91) and novel_circ_002101 (RPM=11422.22) had high expression levels and were specific in Ac1 and Ac2; this suggests that both circRNAs might play special roles in the different developmental stages of the midguts of *A. c. cerana* workers.

Exonic-intronic or exonic circRNAs can regulate the transcription of their source genes by interacting with RNA polymerase II, U1 small nuclear ribonucleoprotein and the gene promoters [40]. In the current work, the GO classifications for the total circRNAs showed that 89 source genes were classified to be associated with metabolism, 36 source genes were classified to be associated with the stimulus response process, and one source gene was classified to be associated with immune system processes; in addition, 130, 129, 112, 112, and 71 source genes were involved in the following categories, respectively: cell process, single tissue process, cell, organelle, and with development. Moreover, the pathway analysis of total circRNAs demonstrated that 89 source genes were annotated to 74 metabolism-related pathways (lipid metabolism, carbohydrate metabolism and energy metabolism, etc.); 126 source genes were annotated to nine immunity-related pathways (JAK-STAT signaling pathway, endocytosis, lysosome, etc.). Additionally, 35, 32, 15, 12, 12 and 10 source genes were annotated with the following development-related signaling pathways: Wnt, Hippo, TGF-β, Hedgehog, mTOR and Notch, respectively. The Wnt signaling pathway is closely related to physiological processes such as mammalian embryo formation, ovary development, and plane cell polarization [44]. It can also affect pigmentation and the development of insect segments, appendages, wings and other organs via cross-talk with the Hippo, Notch and TGF-β signaling pathways [45]. Together, these results indicate that circRNAs are likely to play comprehensive roles in the development of an *A. c. cerana* worker’s midgut.

CircRNAs with differential expression levels were believed to be directly involved in the developmental process of the worker’s midgut. In this study, 34 upregulated circRNAs and 21 downregulated circRNAs were identified in the Ac1 vs Ac2 comparison group, respectively, and their differential expression levels varied widely. Additionally, the differential expression levels of four randomly selected DEcircRNAs were successfully validated using RT-qPCR. The GO analysis showed that an individual DEcircRNA source gene was involved for each of the following processes: cell, cell part, cellular process and organelle; in addition, five and one source genes were involved in catalytic activity and metabolic processes, respectively. Furthermore, the pathway analysis demonstrated that three, two and one source genes were respectively enriched in carbohydrate metabolism, lipid metabolism and nucleotide metabolism. These results indicated that the corresponding DEcircRNA may affect the growth, the development, and the material and energy metabolism of the worker’s midgut by regulating the expression of the source genes. The Hippo signaling pathway regulates organ size by inhibiting cell proliferation and by promoting apoptosis [46], and this pathway interacts with other signaling pathways to regulate gut homeostasis [47]; it also plays a key role in the maintenance of gut structure, especially in the differentiation process of epithelial cells [44]. Here, one source gene of DEcircRNA was enriched in the Hippo signaling pathway. Previous studies suggested that the source genes of DEcircRNAs were significantly enriched in the Hippo signaling pathway during the development of the *B. mori* midgut and the *A. m. ligustica* ovaries [26, 30]. Hence, these results demonstrated that the corresponding DEcircRNAs might play vital roles in the development of insects, including honeybees, via the regulation of the Hippo signaling pathway by controlling their related source genes.

For the honeybee, when an invasion is detected in the hemolymph, the cellular immune responses begin immediately, while after the invasion has continued for many hours, the humoral immune system is activated to induce antimicrobial peptides (AMPs) and bacteriolytic enzymes, and it activates the prophenoloxidase system to defend the body against the invading pathogens [48–50]. Ubiquitin-mediated proteolysis is of great significance for the clearance of senescent and damaged cells [51]. In our study, one and one source genes of DEcircRNAs were enriched in immunity-related pathways such as ubiquitin-mediated proteolysis and lysosome. The MAPK signaling pathway plays an important role in enhancing lymphocyte activity and for the production of immune effector factors that defend against invading pathogens [53]. Here, we found that the source gene of novel_circ_017486, which encodes α-(1,3)-fucosyltransferase C (ncbi_108002060), was associated with antigen expression; in addition, we found that the source gene of novel_circ_009836, which encodes a serine/threonine protein kinase (ncbi_107998746), was associated with the MAPK signaling pathway. These results indicate that DEcircRNAs are able to simultaneously participate in the regulation of both cellular and humoral immunity during the development of the *A. c. cerana* worker’s midgut.

Currently, the functional research of circRNAs is in its infancy, and the function of most circRNAs is still unclear. Salmena et al. [41] proposed the “ceRNA” hypothesis that states that any RNA containing miRNA response elements, such as mRNA, pseudogenes, lncRNA and circRNA, can competitively bind to miRNA. Since then, this hypothesis has been proven by increasing bioinformatic and experimental evidence [27, 29, 54, 55]. For example, by performing anti-AGO2 RNA precipitation and luciferase reporter assays, Ma and colleagues found that circRNA-000284 was closely associated with miR-506 in cervical cancer cells; they also found that the tumor-promoting effect of circRNA-000284 could be abolished by the coexpression of miR-506 mimics or a Snail-2 silencing vector [54]. Recently, by conducting rescue experiments, Zhu et al. [55] observed that circ_0067934 can enhance the proliferation, migration and invasion of hepatocellular carcinoma cells by concomitantly inhibiting miR-1324 and by activating the FZD5/Wnt/β-catenin signaling pathway. In the present work, the regulatory networks of total circRNAs in the worker’s midgut were constructed and analyzed, and the results indicated that 1060 circRNAs could target 74 miRNAs. Among them, most circRNAs (71.51%) had only one target, but a small number of circRNAs can target multiple miRNAs, such as novel_circ_004310 (8 miRNAs), novel_circ_004312 (8 miRNAs), novel_circ_012714 (8 miRNAs), and novel_circ_012715 (8 miRNAs). As shown in the **Fig. 7a**, one miRNA can simultaneously target several circRNAs. For example, miR-6873-y, miR-1895-y and miR-265-y can target 328, 294, 146, and 100 circRNAs, respectively. Moreover, the expression of 13 target miRNAs in the center of the networks was verified by stem-loop RT-PCR (**Fig. 7b**), which further supported the existence of circRNA-miRNA networks in the midguts of *A. c. cerana* workers. To further explore the potential roles of DEcircRNAs, the regulatory networks of DEcircRNAs and their target miRNAs, as well as miRNA-targeted mRNAs in the midgut, were constructed. On the basis of the analytical results, 13 downregulated circRNAs were found to bind to eight miRNAs that target 29 mRNAs, while 16 upregulated circRNAs were found bind to nine miRNAs that target 29 mRNAs. The above results represent a preliminary demonstration that circRNAs acting as ceRNAs could reduce the inhibition or degradation of mRNAs by adsorbing miRNAs, thus affecting the development of the *A. c. cerana* worker’s midgut. Liu et al. [56] discovered that overexpressing miR-184 can promote proliferation but can inhibit the differentiation of adult neural stem/progenitor cells (aNSCs) and can lead to aNSC defects; these defects can be rescued by exogenous *Numbike*, a known regulator of brain development. Here, based on the miRBase database, miR-184-y is highly homologous to miR-184 [57]; this suggests that the circRNAs (novel_circ_012791, novel_circ_012795 and novel_circ_012798, etc.) that target miR-184-y may regulate the proliferation and differentiation of midgut stem cells in an *A. c. cerana* worker. In *Drosophila*, miR-278 mutants were detected to elevate insulin production and the circulating sugar mobilized from adipose-tissue glycogen stores; miR-278 was responsible for maintaining energy homeostasis through the regulation of expanded transcripts [58]. In our study, miR-278-y is highly homologous to miR-278, and it was speculated that miR-278-y, and its corresponding target DEcircRNAs, probably controls the digestion and absorption of the midgut, thus affecting the energy balance in the *A. c. cerana* worker. Ke and colleagues found that the overexpression of miR-149 or the knockdown of *Forkhead box M1* (*FOXM1*) by shRNA could suppress the epithelial-to-mesenchymal transition (EMT) in non-small-cell lung cancer cells (NSCLC cells), whereas increased *FOXM1* expression (partially due to miR-149 downregulation) might contribute to NSCLC cell metastasis [59]. In this work, miR-149-y is highly homologous to miR-149, and we speculated that miR-149-y-targeted DEcircRNAs are likely to be involved in the immune defense of the midguts of *A. c. cerana* workers. Taken together, circRNAs, acting as ceRNAs, can indirectly participate in cell proliferation, differentiation, digestion, absorption, and immune defense in the *A. c. cerana* worker’s midgut via adsorbing the corresponding target miRNAs. Recently, Macedo et al. [60] used sRNA-seq technology to mine miRNAs during the development of the *A. m. ligustica* worker and of the queen’s ovaries and observed 138 expressed miRNAs, including miR-6001. Collins et al. predicted the targets of Bte-miR-6001-5p and Bte-miR- 6001-3p using bioinformatic software and found that many of these targets were likely to be involved in the development and reproductive differentiation of *Apis* and *Drosophila*, including genes associated with ovary and oocyte development, neurodevelopment, larval development and larval molting [61]. In the current investigation, the DEcircRNA-miR-6001-y-mRNA regulatory networks were further constructed and analyzed, and eight DEcircRNAs (novel_circ_005878, novel_circ_018262, novel_circ_005639, novel_circ_015634, novel_circ_004722, novel_circ_016682, novel_circ_015008 and novel_circ_011296) were found to target the same miRNA, miR-6001-y. We inferred that the corresponding DEcircRNAs can affect the development of the midgut by regulating the expression of development-related genes by competitively absorbing miR-6001-y. The aforementioned eight DEcircRNAs can be used as potential candidates for the further study of the molecular mechanisms underlying midgut development.

## Conclusion

In summary, for the first time, circRNAs and corresponding regulatory networks in the midgut of *A. c. cerana* worker were predicted, analyzed and experimentally validated in the current work. Addionally, DEcircRNAs and corresponding ceRNA networks were further explored followed by investigation of potential roles of DEcircRNAs during the development of worker’s midgut.Taken together, these results demonstrated that circRNAs were likely to play a comprehensive part in regulation of growth, development, material and energy metabolism of worker’s midgut; DEcircRNAs might be vital participants in controlling the developmental process of the midgut via regulating the source genes or interacting with target miRNAs as ceRNAs. Our findings can not only provide invaluable resources for future study on *A. c. cerana* circRNA, but also lay a foundation for functional research of key circRNAs involved in the development of eastern honeybee’s midgut.

## Methods

### Honeybee rearing and sample collection

The *A. c. cerana* workers used in this study were obtained from colonies located in the teaching apiary at the College of Bee Science, Fujian Agriculture and Forestry University, Fuzhou city, China.

Sealed brood frames were selected from a healthy colony of *A. c. cerana* and were kept in an incubator at 34 ± 0.5 °C to produce newly emerged honeybees. The emerged workers (termed as 0 d) were carefully removed, confined to plastic cages with holes in groups of 30, kept in the incubator, and fed *ad libitum* with a solution of sucrose (50% w/w in sterile water) using a feeder for 24 h. The feeder was replaced with a new one every 24 h.

Each cage was checked daily, and any dead bees were removed. Ten bees were sacrificed 8 d (or 11 d) after eclosion, and their midguts were quickly processed, frozen in liquid nitrogen and stored at −80 °C for sequencing and molecular experiments. There were three biological replicate cages for each group in this study. The midgut samples collected 8 d after eclosion were named Ac1-1, Ac1-2 and Ac1-3, and those collected at 11 d after eclosion were named Ac2-1, Ac2-2 and Ac2-3.

### RNA extraction, cDNA library construction and deep sequencing

The total RNA of each midgut sample was extracted, and 1.5 μg of RNA per sample was used as input material for rRNA removal using the Ribo-Zero rRNA Removal Kit (Epicentre). cDNA libraries were generated using the NEBNext^®^ Ultra™ Directional RNA Library Prep Kit for Illumina^®^ (NEB) according to the manufacturer’s protocol, and index codes were added to identify each sample. In brief, fragmentation was conducted using divalent cations under elevated temperature in NEBNext First-Strand Synthesis Reaction Buffer (5x). The first-strand cDNA was synthesized using random hexamer primers and reverse transcription. Subsequently, second-strand cDNA synthesis was carried out with DNA Polymerase I and RNase H. Any remaining overhangs were converted into blunt ends via exonuclease/polymerase activities. The NEBNext adaptor with hairpin loop structures was ligated to prepare for hybridization after adenylation of the 3’ ends of the DNA fragments. To select the insert fragments of 150-200□bp in length, the library fragments were purified with AMPure XP Beads (Beckman Coulter). Then, 3□μf USER Enzyme (NEB) was then used with the size-selected, adaptor-ligated cDNA at 37□°C for 15□min. This was followed by PCR amplification with Phusion High-Fidelity DNA polymerase, universal PCR primers and an index (X) primer. Finally, the PCR products were purified (AMPure XP system), and the library quality was assessed using an Agilent Bioanalyzer 2100 and real-time PCR.

The clustering of the index-coded samples was conducted on an acBot Cluster Generation System using TruSeq PE Cluster Kitv3-cBot-HS (Illumina) following the manufacturer’s instructions. After cluster generation, the cDNA libraries were sequenced on an Illumina HiSeq™ 4000 platform (GENE DENOVO Biotechnologies).

### Identification and analysis of circRNA

To identify the circRNAs, we first mapped the clean reads to the *Apis cerana* genome (ACSNU-2.0) [32] using TopHat software [62]. Then, 20 nt from the 5’ and 3’ ends of unmapped reads were extracted and aligned independently to reference sequences by Bowtie2 [63]. Finally, the unmapped anchor reads were submitted to find_circ [8] for the detection of circRNA. Candidate circRNAs were filtered with the following criteria: (1) breakpoints=1; (2) anchor overlap ≤ 2; (3) edit ≤ 2; (4) n uniq > 2; (5) best qual A > 35 or best qual B > 35; (6) n uniq > int (samples/2); (7) circRNA length < 100 kb. The workflow is shown in Additional file 1.

Venn analysis of circRNAs in Ac1 and Ac2 was conducted using the OmicShare online tools (http://www.omicshare.com/tools). The length distribution of circRNAs was calculated and shown in a bar graph using Graph Prism 6 software (GraphPad). Based on the information of circRNAs and the transcripts from the reference genome annotation, the circularization types of circRNA were divided into six groups as follows: annotated exonic circRNA, antisense circRNA, exonic and intronic circRNA, single exonic circRNA, intronic circRNA, and intergenic circRNA.

### Differential expression of circRNA

The expression value of circRNA was normalized to the mapped back-splicing junction reads per million mapped reads (RPM) value. DEcircRNAs were identified using DESeq software [64]. |FC (fold change)| ≥ 2, *p* value < 0.05, and FDR (false discovery rate) ≤ 1 was set as the threshold for significant differential expression by default. The expression clustering of DEcircRNAs was performed using the Omicshare online tools (http://www.omicshare.com/tools).

### Prediction and analysis of source genes

Gene Ontology (GO) and pathway analyses were performed for the source genes of the total predicted circRNAs and the DEcircRNAs. In detail, GO term analysis was performed for the circRNAs using a DAVID gene annotation tool (http://david.abcc.ncifcrf.gov/) [65]. A two-sided Fisher’s exact test was used to classify the GO category, while the false discovery rate (FDR) was calculated to correct the *P* value [66]. The GO terms with a *P* value < 0.05 were considered to be statistically significant. Similarly, a pathway analysis uncovered the significant pathways related to DEcircRNAs according to the annotation of the Kyoto Encyclopedia of Genes and Genomes (KEGG) database (http://www.genome.jp/kegg/) [67]. The significance threshold was defined by a *Q* value (<0.05).

To study the ceRNA roles of circRNA, we predicted the binding pairs of circRNA-miRNA and miRNA-mRNA using TargetFinder [68]. We constructed the regulatory networks of circRNA-miRNA with the top 300 ranked by the *P* values of the hypergeometric distribution and the circRNA-miRNA-mRNA. The regulatory networks were visualized by Cytoscape v.3.2.1 software [69].

### PCR confirmation of novel circRNA with convergent and divergent primers

To verify the back-splicing junction of circRNA, five circRNAs were randomly selected for PCR verification. Special divergent and convergent primers for each candidate circRNA were designed using DNAMAN (Lynnon Biosoft) software following previously described method [21] (Additional file 2: **Table S1**). Total RNA of Ac1 and Ac2 were isolated with AxyPre RNA extraction Kit (Axygen) and divided into three portions, one of which was treated with 3 U/mg RNase R (Geneseed) for 15 min at 37 °C to remove linear RNA followed by reverse transcription with hexamer to synthesize the first strand cDNA, another one portion of RNA was reversed transcribed with OligdT primer to synthesize the first strand cDNA, and the third one portion of RNA was used for stem-loop RT-PCR verification of target miRNAs of circRNAs. Genomic DNA (gDNA) of Ac1 and Ac2 were extracted using AxyPre genomic DNA extraction Kit (Axygen). the equimolar mixture of Ac1 and Ac2 cDNA, as well as the equimolar mixture of Ac1 and Ac2 gDNA were used as templates for PCR. The PCR amplification was carried out on a T100 thermo cycler (Bio-Rad) in a 20 μL reaction volumn containing 1 μL template (200 ng/ μL), 10 μL PCR mixture (TaKaRa), 1 μL upstream primer (10 μmol/L), 1 μL downstream primer (10 μmol/L) and 7 μL sterile water. The reaction parameters were: 94 °C for 5 min; followed by 36 cycles of 94 °C for 40 s, an appropriate annealing temperature (according to the melting temperature of the primers) for 30 s, and 72 °C for 30 s; and 72 °C for 5 min. The PCR products were examined on 2% agarose gel electrophoresis with Genecolor (Gene-Bio) staining.

### Stem-loop RT-PCR verification of DEcircRNA-targeted miRNA

The extracted total RNA of Ac1 and Ac2 samples were treated with DNase I (TaKaRa, China) to remove remaining DNA. Following the previously developed method by Liu et al. [70], stem-loop primers, specific forward primers and universal reverse primers (presented in additional file 6: **Table S7**) were designed using DNAMAN software based on the sequences of the randomly selected nine miRNAs within *A. c. cerana* circRNA-miRNA regulatory networks, including miR-3759-y, miR-4968-y, miR-92-x, novel-m0018-3p, miR-1895-y, miR-7847-y, miR-7550-x, miR-669-y, miR-375-y, miR-965-x, miR-449-x, miR-6717-x and miR-466-y. The primers were synthesized by Sangon Biotech Co., Ltd. One microgram of total RNA was reverse transcribed to cDNA using RevertAid First Strand cDNA Synthesis Kit (TaKaRa) and stem-loop primers. The PCR amplification of randomly selected miRNAs was conducted on a T100 thermo cycler (Bio-Rad) using Premix (TaKaRa) under the following conditions: pre-denaturation step at 94 °C for 5 min; 30 amplification cycles of denaturation at 94 °C for 50 s, annealing at 55 °C for 30 s, and elongation at 72 °C for 1 min; followed by a final elongation step at 72 °C for 10 min. The PCR products were detected on 2% agarose gel eletrophoresis with Genecolor (Gene-Bio) staining.

### Real time quantitative PCR validation of DEcircRNA

To verify the reliability of RNA-seq data, we randomly selected 10 circRNAs for Real time quantitative PCR (RT-qPCR). special divergent primers were designed and cDNA was synthesized according to the method [21] above (Additional file 11: **Table S10**). The RT-qPCR system was 20 μL in including template one μL (200 ng/μL), SYBR Green Dye 10 μL, upstream primer one μL (10 μmol/L), downstream primer one μL (10 μmol/L) and diethyl pyrocarbonate (DEPC) water seven μL. qPCR was performed on ABI 7500 Real time PCR Detection System (ABI, USA). reaction conditon were: 95 °C for 5min; followed by 45 cycles of 94 °C for 15 s, an appropriate annealing temprature (according to the melting temperature of the primers) for 30 s; and then 72 °C for 45s. The *A. c. cerana* housekeeping gene *actin* was used as the internal control. The data were analyzed using 2^−ΔΔ^Ct^^ method and presented as relative expression levels from three biological replicates and three parallel replicates. The data were visualized using Graph Prism 6 software (Graphpad).

### Statistical analysis

Statistical analyses were performed using SPSS 16.0 (IBM) and GraphPad Prism 6.0 software (GraphPad). Data were presented as mean **±** standard deviation (SD). Statistical analysis was calculated using independent-samples t-test and one way ANOVA. Fisher’s exact test was employed to filter the significant GO terms and KEGG pathways using R software 3.3.1 (R Development Core Team). *P* < 0.05 was considered statistically significant.

## Supporting information

Supplemental Figture 1

## Availability of data and materials

The datasets supporting the conclusions of this article are available in the National Centre for Biotechnology Information (NCBI) as BioProject PRJNA487111.

## Authors’ contributions

RG designed this study. DFC, HZC, YD, YLG, CLX, YZZ and CSH carried out laboratory work. RG, DFC, and QYD performed bioinformatic analyses. RG, DFC and HZC supervised the work and contributed to preparation of the manuscript. All authors read and approved the final manuscript.

## Additional file

Additional file 1: **Fig. S1** Pearson correlations between different biological repeats within Ac1 and Ac2 sample groups. **a** Pearson correlations between different biological repeats of Ac1 group. **b** Pearson correlations between different biological repeats of Ac2 group.

Additional file 2: **Table S1** Divergent and convergent primers for RT-PCR assay in this study.

Additional file 3: **Table S2** Summary of GO classifications of source genes of total circRNAs in the *A. c. cerana* worker’s midgut.

Additional file 4: **Table S3** Summary of GO classifications of source genes of DEcircRNAs in Ac1 vs Ac2 comparison group.

Additional file 5: **Table S4** Summary of KEGG database annotations of source genes of total circRNAs in Ac1 and Ac2 comparison group.

Additional file 6: **Table S5** Summary of KEGG database annotations of source genes of DEcircRNAs in Ac1 vs Ac2 comparison group.

Additional file 7: **Table S6** Summary of *A. c. cerana* circRNA-miRNA pairs.

Additional file 8: **Table S7** Primers for stem-loop RT-PCR assay in this study.

Additional file 9: **Table S8** Summary of DEcircRNA-miRNA pairs.

Additional file 10: **Table S9** Summary of target mRNAs of DEcircRNA-targeted miRNAs. Additional file 11: **Table S10** Divergent primers for RT-qPCR assay in this study.

## References

1. Jeck WR, Sorrentino JA, Wang K, Slevin MK, Burd CE, Liu J, et al. Circular RNAs are abundant, conserved, and associated with ALU repeats. RNA. 2013;19:141–157.

2. Lasda E, Parker R. Circular RNAs: diversity of form and function. RNA. 2014;20:1829–1842.

3. He J, Xie Q, Xu H, Li J, Li Y. Circular RNAs and cancer. Cancer Lett. 2017;396:138–144.

4. Qu S, Yang X, Li X, Wang J, Gao Y, Shang R, et al. Circular RNA: A new star of noncoding RNAs. Cancer Lett. 2015;365:141–148.

5. Ashwal-Fluss R, Meyer M, Pamudurti NR, Ivanov A, Bartok O, Hanan M, et al. CircRNA biogenesis competes with pre-mRNA splicing. Mol Cell. 2014;56:55–66.

6. Legnini I, Di Timoteo G, Rossi F, Morlando M, Briganti F, Sthandier O, et al. Circ-ZNF609 is a Circular RNA that can be translated and functions in myogenesis. Mol Cell. 2017;66:22–37.e29.

7. Perkel JM. Assume nothing: the tale of circular RNA. Biotechniques. 2013;55:55–57.

8. Memczak S, Jens M, Elefsinioti A, Torti F, Krueger J, Rybak A, et al. Circular RNAs are a large class of animal RNAs with regulatory potency. Nature. 2013;495:333–338.

9. Salzman J, Chen RE, Olsen MN, Wang PL, Brown PO. Cell-type specific features of circular RNA expression. PLoS Genet. 2013;9:e1003777.

10. Fan X, Zhang X, Wu X, Guo H, Hu Y, Tang F, et al. Single-cell RNA-seq transcriptome analysis of linear and circular RNAs in mouse preimplantation embryos. Genome Biol. 2015;16:148.

11. Zhang C, Wu H, Wang Y, Zhu S, Liu J, Fang X, et al. Circular RNA of cattle casein genes are highly expressed in bovine mammary gland. J Dairy Sci. 2016;99:4750–4760.

12. He L, Zhang A, Xiong L, Li Y, Huang R, Liao L, et al. Deep Circular RNA Sequencing Provides Insights into the Mechanism Underlying Grass Carp Reovirus Infection. Int J Mol Sci. 2017;18.

13. Shen Y, Guo X, Wang W. Identification and characterization of circular RNAs in zebrafish. FEBS Lett. 2017;591:213–220.

14. Zhang XH, Yan YM, Lei XY, Li AJ, Zhang HM, Dai ZK, et al. Circular RNA alterations are involved in resistance to avian leukosis virus subgroup-J-induced tumor formation in chickens. Oncotarget. 2017;8:34961–34970.

15. Huang M, Shen Y, Mao H, Chen L, Chen J, Guo X, et al. Circular RNA expression profiles in the porcine liver of two distinct phenotype pig breeds. Asian-Australas J Anim Sci. 2018;31:812–819.

16. Lu T, Cui L, Zhou Y, Zhu C, Fan D, Gong H, et al. Transcriptome-wide investigation of circular RNAs in rice. RNA. 2015;21:2076–2087.

17. Darbani B, Noeparvar S, Borg S. Identification of Circular RNAs from the parental genes involved in multiple aspects of cellular metabolism in barley. Front Plant Sci. 2016;7:776.

18. Sun X, Wang L, Ding J, Wang Y, Wang J, Zhang X, et al. Integrative analysis of *Arabidopsis thaliana* transcriptomics reveals intuitive splicing mechanism for circular RNA. FEBS Lett. 2016;590:3510–3516.

19. Danan M, Schwartz S, Edelheit S, Sorek R. Transcriptome-wide discovery of circular RNAs in Archaea. Nucleic Acids Res. 2012;40:3131–3142.

20. Cortes-Lopez M, Gruner MR, Cooper DA, Gruner HN, Voda AI, van der Linden AM, et al. Global accumulation of circRNAs during aging in *Caenorhabditis elegans*. BMC Genomics. 2018;19:8.

21. Guo R, Chen D, Chen H, Fu Z, Xiong C, Hou C, et al. Systematic investigation of circular RNAs in *Ascosphaera apis*, a fungal pathogen of honeybee larvae. Gene. 2018.

22. Guo R, Chen D, Chen H, Xiong C, Zheng Y, Hou C, et al. Genome-wide identification of circular RNAs in fungal parasite *Nosema ceranae*. Curr Microbiol. 2018.

23. Toptan T, Abere B, Nalesnik MA, Swerdlow SH, Ranganathan S, Lee N, et al. Circular DNA tumor viruses make circular RNAs. Proc Natl Acad Sci U S A. 2018.

24. Westholm JO, Miura P, Olson S, Shenker S, Joseph B, Sanfilippo P, et al. Genome-wide analysis of *drosophila* circular RNAs reveals their structural and sequence properties and age-dependent neural accumulation. Cell Rep. 2014;9:1966–1980.

25. Gan H, Feng T, Wu Y, Liu C, Xia Q, Cheng T. Identification of circular RNA in the *Bombyx mori* silk gland. Insect Biochem Mol Biol. 2017;89:97–106.

26. Chen X, Shi W, Chen C. Differential circular RNAs expression in ovary during oviposition in honey bees. Genomics. 2018.

27. Du WW, Yang W, Liu E, Yang Z, Dhaliwal P, Yang BB. Foxo3 circular RNA retards cell cycle progression via forming ternary complexes with p21 and CDK2. Nucleic Acids Res. 2016;44:2846–2858.

28. Yang L, Han B, Zhang Y, Bai Y, Chao J, Hu G, et al. Engagement of circular RNA HECW2 in the nonautophagic role of ATG5 implicated in the endothelial-mesenchymal transition. Autophagy. 2018;14:404–418.

29. Cheng X, Zhang L, Zhang K, Zhang G, Hu Y, Sun X, et al. Circular RNA VMA21 protects against intervertebral disc degeneration through targeting miR-200c and X linked inhibitor-of-apoptosis protein. Ann Rheum Dis. 2018;77:770–779.

30. Hu X, Zhu M, Liu B, Liang Z, Huang L, Xu J, et al. Circular RNA alterations in the *Bombyx mori* midgut following *B. mori* nucleopolyhedrovirus infection. Mol Immunol. 2018;101:461–470.

31. Hora ZA, Altaye SZ, Wubie AJ, Li J. Proteomics improves the new understanding of honeybee biology. J Agr Food Chem. 2018;66:3605–3615.

32. Park D, Jung JW, Choi BS, Jayakodi M, Lee J, Lim J, et al. Uncovering the novel characteristics of Asian honey bee, *Apis cerana*, by whole genome sequencing. BMC Genomics. 2015;16:1.

33. Tan K, Dong S, Li X, Liu X, Wang C, Li J, et al. Honey bee inhibitory signaling is tuned to threat severity and can act as a colony alarm signal. PLoS Biol. 2016;14:e1002423.

34. Diao Q, Sun L, Zheng H, Zeng Z, Wang S, Xu S, et al. Genomic and transcriptomic analysis of the Asian honeybee *Apis cerana* provides novel insights into honeybee biology. Sci Rep. 2018;8:822.

35. Li HL, Zhang YL, Gao QK, Cheng JA, Lou BG. Molecular identification of cDNA, immunolocalization, and expression of a putative odorant-binding protein from an Asian honey bee, *Apis cerana cerana*. J Chem Ecol. 2008;34:1593–1601.

36. Nishiura JT, Burgos C, Aya S, Goryacheva Y, Lo W. Modulation of larval nutrition affects midgut neutral lipid storage and temporal pattern of transcription factor expression during mosquito metamorphosis. J Insect Physiol. 2007;53:47–58.

37. Kumar S, Molina-Cruz A, Gupta L, Rodrigues J, Barillas-Mury C. A peroxidase/dual oxidase system modulates midgut epithelial immunity in *Anopheles gambiae*. Science. 2010;327:1644–1648.

38. Ellegaard KM, Tamarit D, Javelind E, Olofsson TC, Andersson SG, Vasquez A. Extensive intra-phylotype diversity in lactobacilli and bifidobacteria from the honeybee gut. BMC Genomics. 2015;16:284.

39. Kwong WK, Moran NA. Evolution of host specialization in gut microbes: the bee gut as a model. Gut microbes. 2015;6:214–220.

40. Li Z, Huang C, Bao C, Chen L, Lin M, Wang X, et al. Exon-intron circular RNAs regulate transcription in the nucleus. Nat Struct Mol Biol. 2015;22:256–264.

41. Salmena L, Poliseno L, Tay Y, Kats L, Pandolfi PP. A ceRNA hypothesis: the Rosetta Stone of a hidden RNA language? Cell. 2011;146:353–358.

42. Hansen TB, Jensen TI, Clausen BH, Bramsen JB, Finsen B, Damgaard CK, et al. Natural RNA circles function as efficient microRNA sponges. Nature. 2013;495:384–388.

43. Li RC, Ke S, Meng FK, Lu J, Zou XJ, He ZG, et al. CiRS-7 promotes growth and metastasis of esophageal squamous cell carcinoma via regulation of miR-7/HOXB13. Cell death & disease. 2018;9:838.

44. Fevr T, Robine S, Louvard D, Huelsken J. Wnt/beta-catenin is essential for intestinal homeostasis and maintenance of intestinal stem cells. Mol Cell Biol. 2007;27:7551–7559.

45. Aronstein KA, Murray KD. Chalkbrood disease in honey bees. J Invertebr Pathol. 2010;103 Suppl 1:S20–29.

46. Halder G, Johnson RL. Hippo signaling: growth control and beyond. Development. 2011;138:9–22.

47. Camargo FD, Gokhale S, Johnnidis JB, Fu D, Bell GW, Jaenisch R, et al. YAP1 increases organ size and expands undifferentiated progenitor cells. Curr Biol. 2007;17:2054–2060.

48. Hedengren-Olcott M, Olcott MC, Mooney DT, Ekengren S, Geller BL, Taylor BJ. Differential activation of the NF-kappaB-like factors Relish and Dif in *Drosophila* melanogaster by fungi and Gram-positive bacteria. J Biol Chem. 2004;279:21121–21127.

49. Evans JD, Aronstein K, Chen YP, Hetru C, Imler JL, Jiang H, et al. Immune pathways and defence mechanisms in honey bees *Apis mellifera*. Insect Mol Biol. 2006;15:645–656.

50. Stanley D, Miller J, Tunaz H. Eicosanoid actions in insect immunity. J Innate Immun. 2009;1:282–290.

51. McBride WH, Iwamoto KS, Syljuasen R, Pervan M, Pajonk F. The role of the ubiquitin/proteasome system in cellular responses to radiation. Oncogene. 2003;22:5755–5773.

52. Chen D, Guo R, Xu X, Xiong C, Liang Q, Zheng Y, et al. Uncovering the immune responses of *Apis mellifera ligustica* larval gut to *Ascosphaera apis* infection utilizing transcriptome sequencing. Gene. 2017;621:40–50.

53. Saraav I, Singh S, Sharma S. Outcome of Mycobacterium tuberculosis and Toll-like receptor interaction: immune response or immune evasion? Immunol Cell Biol. 2014;92:741–746.

54. Ma HB, Yao YN, Yu JJ, Chen XX, Li HF. Extensive profiling of circular RNAs and the potential regulatory role of circRNA-000284 in cell proliferation and invasion of cervical cancer via sponging miR-506. Am J Transl Res. 2018;10:592–604.

55. Zhu Q, Lu G, Luo Z, Gui F, Wu J, Zhang D, et al. CircRNA circ_0067934 promotes tumor growth and metastasis in hepatocellular carcinoma through regulation of miR-1324/FZD5/Wnt/beta-catenin axis. Biochem Biophys Res Commun. 2018;497:626–632.

56. Liu C, Teng ZQ, Santistevan NJ, Szulwach KE, Guo W, Jin P, et al. Epigenetic regulation of miR-184 by MBD1 governs neural stem cell proliferation and differentiation. Cell Stem Cell. 2010;6:433–444.

57. Griffiths-Jones S. miRBase: microRNA sequences and annotation. Current protocols in bioinformatics. 2010;12:11–10.

58. Teleman AA, Maitra S, Cohen SM. *Drosophila* lacking microRNA miR-278 are defective in energy homeostasis. Genes Dev. 2006;20:417–422.

59. Ke Y, Zhao W, Xiong J, Cao R. miR-149 inhibits non-small-cell lung cancer cells EMT by targeting FOXM1. Biochem Res Int. 2013;2013:506731.

60. Macedo LM, Nunes FM, Freitas FC, Pires CV, Tanaka ED, Martins JR, et al. MicroRNA signatures characterizing caste-independent ovarian activity in queen and worker honeybees *(Apis mellifera* L.). Insect Mol Biol. 2016;25:216–226.

61. Collins DH, Mohorianu I, Beckers M, Moulton V, Dalmay T, Bourke AF. MicroRNAs associated with caste determination and differentiation in a primitively eusocial Insect. Sci Rep. 2017;7:45674.

62. Kim D, Pertea G, Trapnell C, Pimentel H, Kelley R, Salzberg SL. TopHat2: accurate alignment of transcriptomes in the presence of insertions, deletions and gene fusions. Genome Biol. 2013;14:R36.

63. Langdon WB. Performance of genetic programming optimised Bowtie2 on genome comparison and analytic testing (GCAT) benchmarks. Biodata Min. 2015;8:1.

64. Wang L, Feng Z, Wang X, Wang X, Zhang X. DEGseq: an R package for identifying differentially expressed genes from RNA-seq data. Bioinformatics. 2010;26:136–138.

65. Huang DW, Sherman BT, Tan Q, Collins JR, Alvord WG, Roayaei J, et al. The DAVID gene functional classification tool: a novel biological module-centric algorithm to functionally analyze large gene lists. Genome Biol. 2007;8:R183.

66. Jung SH. Stratified Fisher’s exact test and its sample size calculation. Biom J. 2014;56:129–140.

67. Du J, Yuan Z, Ma Z, Song J, Xie X, Chen Y. KEGG-PATH: Kyoto encyclopedia of genes and genomes-based pathway analysis using a path analysis model. Mol biosyst. 2014;10:2441–2447.

68. Allen E, Xie Z, Gustafson AM, Carrington JC. microRNA-directed phasing during trans-acting siRNA biogenesis in plants. Cell. 2005;121:207–221.

69. Smoot ME, Ono K, Ruscheinski J, Wang PL, Ideker T. Cytoscape 2.8: new features for data integration and network visualization. Bioinformatics. 2011;27:431–432.

70. Liu F, Peng W, Li Z, Li W, Li L, Pan J, et al. Next-generation small RNA sequencing for microRNAs profiling in *Apis mellifera:* comparison between nurses and foragers. Insect Mol Biol. 2012;21:297–303.

