## Supplemental Table 1 for "Systematic identification of circular RNAs and corresponding regulatory networks unveil their potential roles in the midgut of *Apis cerana cerana* workers"

**Table S1** Divergent primers and convergent primers for RT-PCR

| **circRNA** | **Primer** | **Primer sequence (5'-3')** | **Product size** |
| --- | --- | --- | --- |
| novel_circ_012591 | D-F | GAAGAGGATTTGGCAGACC | 124 bp |
|  | D-R | AGAGTTTGTTGTTCCCTTGG |  |
|  | C-F | CCAATCACTTCTACTTCACCG | 129 bp |
|  | C-R | TCTCCGTCGTTAGGTTCAA |  |
| novel_circ_014624 | D-F | ACCAGTATCTTCGCCCTCA | 114 bp |
|  | D-R | GTTTCTTGTCGTTGTCCGA |  |
|  | C-F | CATCAACATCAACATCAGCC | 133 bp |
|  | C-R | GAAGGTTCAACGGAAGATTG |  |
| novel_circ_000659 | D-F | CGAAGCATTGAGGAAGAAG | 121 bp |
|  | D-R | AGTCTCTCGGTCTGGACATC |  |
|  | C-F | TGTCCAGACCGAGAGACTTG | 150 bp |
|  | C-R | GCTGATGATGCTGTTGTAGC |  |
| novel_circ_012668 | D-F | ATGAACTCACGCAAGAGCA | 81 bp |
|  | D-R | ATCGTCTGGATTGGTTGG |  |
|  | C-F | AGAAGATTCCAGTAGCGAGAAG | 134 bp |
|  | C-R | GAGTAGGCAGGGTGATACCTT |  |
| novel_circ_000028 | D-F | CATTGCTCCGACTTGGTCT | 111 bp |
|  | D-R | TGTGGTTGTCTTGGTTGATG |  |
|  | C-F | GTGGGACTCACTGACAAGACT | 117 bp |
|  | C-R | GAGCCTTACTATCACAGATGGG |  |
| *actin* | D-F | GGTTGTTGATAGTGGAGATGG | 198 bp |
|  | D-R | CACGACCAGCAATAGGAAT |  |
|  | C-F | TACAGAAATACGCCAATA | 98 bp |
