## Supplemental Table 10 for "Systematic identification of circular RNAs and corresponding regulatory networks unveil their potential roles in the midgut of *Apis cerana cerana* workers"

**Table S10** Primers for RT-qPCR

| **CircRNA name** | **Primers** | **Primer sequences (5'-3')** |
| --- | --- | --- |
| novel_circ_018262 | D-F | TGCGGTTGCTATGGGATT |
|  | D-R | TCCTCGCCGTTCAAACAT |
| novel_circ_012576 | D-F | CACCATCATCCTCGAAACG |
|  | D-R | TTCTTGGGCCGGGAAAGC |
| novel_circ_011531 | D-F | GGTGTCGTGCGACCAGTT |
|  | D-R | GGGAGCCTGTGGTATTGG |
| novel_circ_002054 | D-F | GGATTCCTACCGCAGATT |
|  | D-R | CGATGTCCAGAGGTCCAG |
| *actin* | C-F | TACAGAAATACGCCAATA |
|  | C-R | TAAAGATAAAGCAGAAGC |
