## Supplementary figures and images for "Systematic identification of circular RNAs and corresponding regulatory networks unveil their potential roles in the midgut of *Apis cerana cerana* workers"

### Supplemental Figture 1

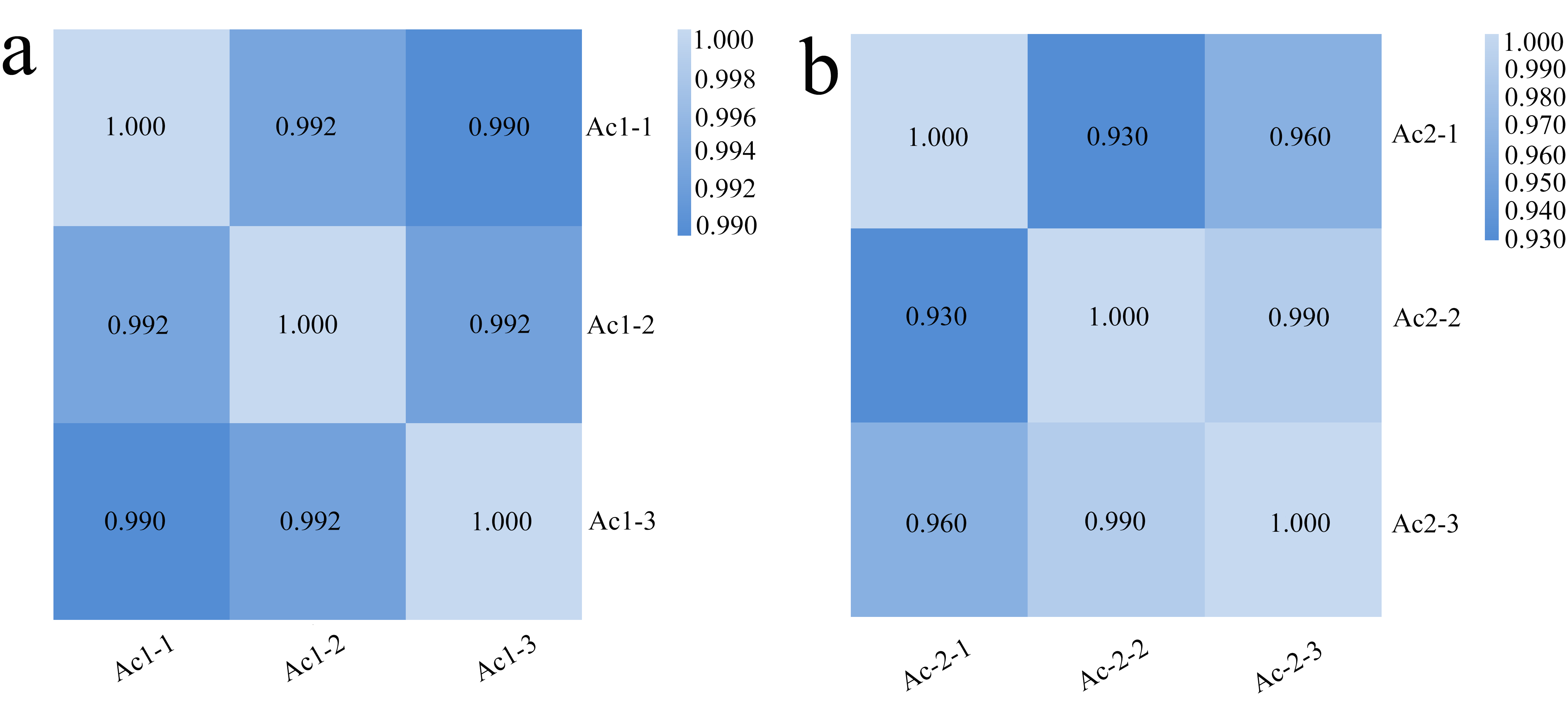
